# High resolution structure of plant Light-Harvesting Complex II (LHCII) provides insight into lutein conformations and energy quenching

**DOI:** 10.64898/2026.07.22.740098

**Authors:** Tasmin Spurgeon, Stephen P. Muench, Peter G. Adams

## Abstract

Plant Light Harvesting Complex II (LHCII) is found in the thylakoid membranes of chloroplasts and balances two roles: energy collection for photosynthesis and energy dissipation to prevent photo-damage when there is excessive sunlight. The mechanism for LHCII to switch between two energetic states has been debated but may involve a pH-triggered conformational change. Here we present single-particle cryo-electron microscopy (EM) structures of “light-harvesting” LHCII in detergent at pH 7.5 and pH 4.5. The high resolution (2.48 Å) maps provide clear placements for all bound pigments, giving high confidence in the models. Surprisingly, we find that there is little conformational change to the polypeptide between these new light harvesting structures and previously published crystals structures, thought to be energy dissipating. The crossing angles of helix A/B and the Lutein 1-Chlorophyll 612 separation distances are similar. This contrasts with other recent analyses of LHCII by single-particle EM that suggested a change to the helix A/B angle and a reduction in Lutein 1-Chlorophyll 612 separation may trigger quenching and a photoprotective state. The high resolution of our structures also allowed us to investigate small conformational changes of the lutein within L1/L2 binding sites of LHCII, revealing rotations and distortions in the pigment that could lead to changes in energy transfer. In addition, we find that low pH causes LHCII to form a destabilised structure where pigment loss from the V1 binding site (usually violaxanthin or zeaxanthin) correlated with a disordered C-terminus, often for just one LHCII monomer with an LHCII trimer. Overall, our findings have important implications for the molecular mechanism of photoprotection.

## Introduction

Light harvesting complex II (LHCII) is a pigment-protein complex found in the thylakoid membrane in chloroplasts of higher plants and is essential for efficient photosynthesis, carrying out multiple roles (Croce & van Amerongen, 2011). LHCII serves as the primary antenna for Photosystem II (PSII), increasing the quantity and wavelengths of light that plants can absorb through the array of pigment it contains (van Amerongen & van Grondelle, 2001). After LHCII absorbs a photon, the resultant excitation energy is passed to PSII for conversion to chemical energy. However, excessive excitation energy risks damaging PSII by causing photo-oxidative reactions, therefore, under high light conditions, the process of non-photochemical quenching (NPQ) is activated (Janik-Zabrotowicz & Gruszeki, 2023; Navakoudis et al., 2023). There are different mechanisms involved in NPQ but the fastest acting, named energy-dependant quenching (qE), involves LHCII (Ruban et al., 2012; Ruban & Wilson, 2021). The pH of the thylakoid lumen drops due to excessive excitation leading to greater electron flow and proton pumping, triggering a pathway which ultimately results in LHCII switching to a photoprotective state. In this quenched state, a larger fraction of the harvested energy is dissipated as heat instead of being transferred to PSII, via a much-debated mechanism. Previous reports have suggested that the fluorescence lifetime of LHCII indicates its different states: the light harvesting state has a lifetime of ∼4ns, while the quenched state is far shorter at <0.5ns (Chmeliov et al., 2019; Gray et al., 2024; Mullineaux et al., 1993). The exact process leading to qE and the mechanism by which LHCII dissipates energy remains unclear. Single-protein fluorescence measurements have shown that LHCII in isolation can spontaneously swap to a quenched state and the proportion can be increased by a large drop in pH, clearly indicating LHCII is the source of energy dissipation (Schlau-Cohen et al., 2015; Schörner et al., 2015). However, the low pH required to activate this quenched state in vitro, around pH 4.5, is significantly lower than the physiologically relevant pH thought to reach as low as of 5.4 in the native environment, therefore, other factors must sensitise LHCII (Johnson & Ruban, 2011; X.-P. Li et al., 2000; Nicol & Croce, 2021; Park et al., 2018; Saccon, Giovagnetti, et al., 2020; Shukla et al., 2020). One such factor is the violaxanthin de-epoxidase enzyme which, under high light intensities, converts the carotenoid (Car) pigment in the V1 binding site of LHCII from violaxanthin (Vio) to zeaxanthin (Zea), making LHCII more sensitive to lowered pH (Sacharz et al., 2017). How the violaxanthin de-epoxidase enzyme accesses this V1-bound pigment remains unclear. PsbS is another factor required for the NPQ response under physiological conditions, a membrane protein that is sensitive to pH and may interact with and promote the state change of LHCII, potentially via a conformational change itself related to hydrophobic mismatch (Correa-Galvis et al., 2016; Krishnan-Schmieden et al., 2021; Liguori et al., 2019; Wilson et al., 2024).

The identity of the quenching pigment and the precise photophysical mechanism of energy dissipation within LHCII has been disputed. Aggregation of LHCII, either in artificial 2-D or 3-D aggregates or within model lipid vesicles, has been shown to reduce the fluorescence lifetime of the system, possibly mimicking the thylakoid membrane (Adams et al., 2018; Liu et al., 2004; van Oort et al., 2007; Wilson et al., 2022). However, individual non-aggregated LHCII can also be observed with a short fluorescence lifetime in a variety of *in vitro* contexts, both isolated in gels and detergent solutions, suggesting there is a mechanism, involving an individual LHCII trimer, that results in an NPQ-like state that can quench excitation energy within LHCII (Krüger et al., 2010; Saccon, Durchan, et al., 2020; Schlau-Cohen et al., 2015). Several predictions have been made regarding the underlying mechanism of qE based on a variety of molecular simulations, energetic calculations and structural information. The majority of studies agree that one of the Cars would be the final quenching site as they have somewhat unique energetics which include a dipole-forbidden dark S_1_ state that has a very short, sub-ps, excited state lifetime that can dissipate energy as heat. Various ultrafast spectroscopy studies have suggested the photophysical mechanism could involve either a hot S_1_, S*/ S_n_, charge-transfer (CT), or hybrid exciton/CT state (Liguori et al., 2017; Macernis et al., 2012; Pedraza-González et al., 2024; Son et al., 2019). Of the four Cars in LHCII, the lutein (Lut) in the L1 site, Lut1, is a likely candidate as it is positioned closely to the lowest energy Chl trio of Chl *a* 610-612, an optimal location to accept and dissipate excitation energy within LHCII (Fox et al., 2017). However, there is still debate over the protein structural change that determines the switch between LHCII passing energy on to PSII (light harvesting state) or dissipating the energy (quenching state). It has been suggested that a conformational change in LHCII, such as the shifting of α-helices, may bring Lut1 and the low energy Chl trio closer, resulting in a change in energetic coupling, ultimately leading to quenching (Daskalakis et al., 2019; H. Li et al., 2020). Other hypotheses involve squeezing LHCII, perhaps by lipid rearrangements, also resulting in similar pigment rearrangements which cause the quenching (Tietz et al., 2020; Wilson et al., 2024). Alternatively, rather than reducing the separation of pigments, a change in the molecular configuration of the Lut1 pigment (e.g., a twist) has been suggested to induce new or modified excitation levels that are below the the lowest-energy Chls, so that Lut1 would be biased towards receiving the excitation energy from the low-energy Chl and dissipating it (Liguori et al., 2015; Ma et al., 2025; Macernis et al., 2012).

The general structure of trimeric LHCII of higher plants has been solved by X-ray crystallography revealing three transmembrane α-helices per monomer (helix A, B and E), an amphiphilic helix (helix D) and a short 3_10_-helix (helix E) with an arrangement of four Cars and 14 Chl (Kühlbrandt et al., 1994; Liu et al., 2004; Standfuss et al., 2005). The highest resolution achieved was 2.72 Å and the resulting LHCII atomic model identified all Chl *a* and Chl *b* (Liu et al., 2004). However, the LHCII within these protein crystals appeared to be in the quenched, qE, state as judged from their short fluorescent lifetimes (van Oort et al., 2011). In recent years, the advent of improved cryogenic electron microscopy (cryoEM) instrumentation and processing methods has allowed a structural model to be calculated for LHCII isolated in detergent micelles or lipid nanodiscs where the protein is expected to be in the light harvesting state from its high fluorescence lifetime. Ruan et al.(2023) reported a new LHCII model that contained rearrangements of pigments and the formation of an additional α-helix at the C-terminus. However, the map quality near the key pigments in the structure was imperfect which leaves ambiguity in the pigment conformations.

In the current work, we have taken advantage of further developments in cryoEM technology to obtain a higher resolution, “light harvesting” structure of LHCII from spinach in a detergent micelle at both pH 7.5 and pH 4.5, which all have long fluorescence lifetimes. The quality of our electron density map allows the building of a structural model of the protein and associated co-factors with high confidence, allowing a direct comparison to previously published short lifetime, crystal structures. Surprisingly, we observe a high degree of similarity between our structures and the crystal structures, particularly in the polypeptide and pigment separation, contrary to some established hypothesis. Instead, we do find clear structural evidence for a Lut1 conformational change. We perform comprehensive measurements of the architecture within LHCII, comparing our structural model to others, and discuss the possible implications of our findings, in the context of a potential NPQ switch.

## Materials and Methods

### Protein isolation and purification

Spinach was purchased on the morning of the purification and kept in the dark at 4 °C for approximately 0.5-1 hr and then trimeric LHCII was extracted and purified following previously published protocol with minor modification (Adams et al., 2018). Briefly, spinach leaves were macerated in ice-cold buffer A (300 mM sucrose, 5 mM EDTA, 50 mM HEPES, pH 7.5), the liquid was filtered through muslin cloth and cottonwool, then chloroplasts were collected by centrifugation, resuspended in buffer B (5 mM EDTA, 10 mM Tricine pH 7.4) and osmotically lyzed by adding an equal volume of buffer & (400 mM sucrose, 5 mM EDTA, 10 mM Tricine, pH 7.4). Isolated thylakoid membranes were adjusted to 0.5 mg Chl/mL and solubilized with 1% (w/v) detergent n-dodecyl α-D-maltoside (α-DDM, Anatrace) in 20 mM HEPES buffer pH 8 for 1 hr on ice. Thylakoid membrane proteins were then separated via ultracentrifugation on sucrose density gradients (8-13% w/w sucrose at 100,000 × g, 36 hr, 4 °C). The LHCII trimer band was then collected, concentrated using 30 kDa Vivaspin Ultra centrifugal filters (Merck Millipore, UK), and further purified using high-resolution size exclusion chromatography in 150 mM NaCl, 0.03% α-DDM, 20 mM HEPES (pH 7.5) using a 16/600 Superdex 200 prep grade column on an AKTA Prime FPLC system (GE Healthcare Life Sciences, PA, USA). After pooling the appropriate eluted fractions, they were concentrated using Vivaspin centrifugal filters once more.

The sample pH was adjusted as required by using a Vivaspin20 100,000 MWCO concentration spin column to exchange the buffer (that contained high NaCl after size-exclusion chromatography) for a new buffer containing 20mM HEPES, 0.03% α-DDM at either pH 7.5 or pH 4.5 (as appropriate for the sample) to give an absorbance of 30-40 at 680 nm. LHCII trimer molar concentration was estimated as [Chl mM concentration] × 42, where “chlorophyll mM concentration” is determined from absorbance measurements after methanol/acetone pigment extraction. Typically, the concentration of LHCII trimers was found to be approx. 100 nM in buffer of 20 mM HEPES (pH 7.5). SDS-PAGE (NuPAGE® Novex® 12% Bis-Tris Protein Gel with Precision Plus Protein Dual Color Standards, Bio Rad ladder) and Native-PAGE (NativePAGE™ Novex® 4-16% Bis-Tris Protein Gels) was used to confirm protein purity and oligomerisation state.

### Absorbance and fluorescence spectroscopy

UV-Vis absorption spectra were collected using an Agilent Cary 5000 spectrophotometer with samples diluted in working buffer (20mM HEPES, 0.03% α-DDM, pH 7.5 or 4.5) to give an absorbance of approximately 0.1 at 680 nm. Steady state emission spectra and fluorescence decay were acquired with an Edinburgh Instruments FLS 980 fluorimeter. Fluorescence decay data was fitted using F980 software (Edinburgh Instruments). All spectroscopy measurements were taken with a PMMA (UV-transparent) plastic cuvette of 10 mm pathlength.

### CryoEM Data Collection

The purified LHCII was adjusted to the appropriate pH (absorbance of 30-40 at 680 nm). For each sample, 3 μL was applied to a holey carbon grid (Cu, 300 mesh, 1.2/1.3, Quantifoil) after glow discharge (45 s, 0.38 mBar, PELCO easiGlow). Grids were blotted and plunge frozen in liquid ethane (Vitribot Mark II). Micrographs were collected on a 300 kV Titan Krios in counting mode using EPU 3.3 software (Thermo Fisher Scientific). See Table S1 for all parameters.

### EM data processing

All processing was carried out using CryoSparc v4.7.1 (Punjani et al. 2017). Micrographs were pre-processed with “Patch Motion correction” and “Patch CTF Estimation”. Blob picker was used to auto-select particles on denoised micrographs to selected 2,786,533 and 2,774,012 particles for the pH 7.5 and 4.5 data sets, respectively. Two rounds of 2-D classification were used to remove junk particles resulting in particle stacks of 208,480 and 354,974. Initial models were made using “ab-initio reconstruction” with five classes followed by heterogenous reconstruction.

After trial and error one model and associated particles were taken forward for the pH 7.5 data set while two were taken for the pH 4.5 set. It was determined the remaining pH 7.5 particles were either junk or of poorer quality, from regions of thicker ice, and did not represent any alternative conformation. Meanwhile, the pH 4.5 data set contained two clear distributions of particles. With the resultant particle stack, non-uniform refinement was carried out for each data set followed by re-extraction with a larger 364 px box size and “Local CTF refinement”, “Global CTF refinement” and “Reference based motion correction” was carried out (Punjani et al., 2020; Rubinstein & Brubaker, 2015; Zivanov et al., 2019, 2020). The pH 7.5 data set then underwent a final non-uniform refinement to achieve the final map at 2.48 Å. Various methods were attempted to identify any alternative conformation in the pH 7.5 data set, included the one outlined below used for the pH 4.5 data set, but no alternative state was found.

The pH 4.5 data set was determined to have mixed states due to partial density, specifically around the V1 binding sites and C-terminus, therefore, different approaches were tried to separate these states. Ultimately, 3-D classification, with a focus mask over the V1 site, was used to produce five classes. One class contained particles with fully occupied V1 site and was used for non-uniform refinement to yield the “pH 4.5” map at 2.60 Å from 40,661 particles. The other four classes had a mix of occupied, partially occupied and empty V1 sites, where the site was empty the C-terminus was disordered. One of these classes had a monomer with an unoccupied V1 site, another with a partial occupancy and the final monomer had an occupied V1 binding site and this was picked as representative of a destabilised state. Non-uniform refinement with this map and particle stack of 39,449 particles was used to construct the final “destabilised pH 4.5” map at 2.68 Å resolution. The workflow for both data sets can be seen in Fig. S1.

C1 symmetry (no symmetry at all) was used throughout for all reconstructions. Local resolution was estimated with the “local resolution estimation” job and the final resolutions determined by gold standard FSC. All 3-D structural visualisations were done with ChimeraX.

### Real-space refinement of EM data

The PDB 8IWX (Ruan et al., 2023) was used as a starting model, and rigid body fitting was used to fit it into the pH 7.5 map before real space refinement was carried out in Win*Coot* (Emsley et al., 2010). The resultant model was used in the same way to model the two pH 4.5 models. Isolde (Croll, 2018) was used to refine the Lut molecules in all models and the C-terminus of the destabilised pH 4.5 model, though there is minimal density for the latter.

### Measurements of distances and angles within atomic models of LHCII structure

ChimeraX was used to visually compare all maps and models and to calculate measurements to quantify differences. Fig. S2B defines each measurement used in this paper. Distances were measured using the distance command specifying two atoms; twists and rotations were measured using the torsion command specifying four atoms. For each structure, measurements were taken from the three monomers, averaged and a standard deviation calculated. In our LHCII models, each monomer was independently modelled, therefore, the standard deviation calculated is a cumulation of the natural variation present within trimers, and the variations caused by modelling differences. In some other LHCII structural models where three-fold symmetry was applied, the standard deviation will instead represent the variation in modelling.

## Results

### Protein production and function

The LHCII protein was extracted from spinach leaves and purified as described in the Methods, following an established procedure (Adams et al., 2018; Hancock et al., 2022). The sucrose gradient band pattern of the separated thylakoid membrane can be seen in Fig. 1A: the upper light-green band contains monomeric LHCII while the lower dark-green band contains trimeric LHCII, consistent with previous work. SDS-PAGE and Native-PAGE electrophoresis gels were used to confirm the identity and purity of the protein after sucrose gradient purification and we found that both samples’ purity was further improved by size exclusion chromatography (SEC), as evidenced by the fading of the additional bands in the SDS-PAGE gels (Fig. 1B). LHCII monomers were successfully removed by SEC with a single band present in the Native-PAGE gel (Fig. 1C) that represents LHCII trimers (by comparison with previous work). LHCII samples were diluted in a working buffer (20mM HEPES, 0.03% α-DDM) adjusted to either pH 7.5 or 4.5 and characterised via absorption and fluorescence spectroscopy. The absorption spectra show the expected pattern for LHCII trimers: a Chl *a* Q_y_ maximum at 675 nm and the expected peaks between 420 nm and 490 nm, including a shoulder around 480 nm, are indicative of intact trimeric LHCII (Fig. 1D, solid lines) (Peterman et al., 1997). The single peak at 681 nm in the fluorescence emission spectrum, also supports energetic connectivity of the pigment network within LHCII (Fig. 1D, dashed lines). Fluorescence decay curves were measured and fitted to a tri-exponential decay function revealing a mean fluorescence lifetime of ∼4.2 ns for the sample at pH 7.5 and ∼4.3 ns for the sample at pH 4.5, values that are within error of each other and consistent with a light harvesting state of LHCII (Fig. 1E). As it has been previously reported that LHCII requires a large drop in pH to initiate a change to energy dissipating states (in the absence of PsbS and Zea) (Johnson & Ruban, 2011), we characterised the extracted LHCII at pH 7.5, 5.4, 4.5 and 3.5 to determine if the fluorescence lifetime would reduce. Regardless of pH, no significant reduction in fluorescence lifetime was observed in these samples; the only change was a small reduction in fluorescence intensity at pH 3.5 which may indicate damage to the protein (Fig. S3). Ultimately, pH 7.5 was chosen as the standard conditions for the first LHCII sample to represent the light-harvesting state, and pH 4.5 was selected as the conditions for a second LHCII sample where this reduced pH may yet change the protein conformation (despite the similar fluorescence lifetime), chosen as the lowest pH that did not significantly change the emission spectrum (Do et al., 2026; Johnson & Ruban, 2011).

**Fig. 1.**
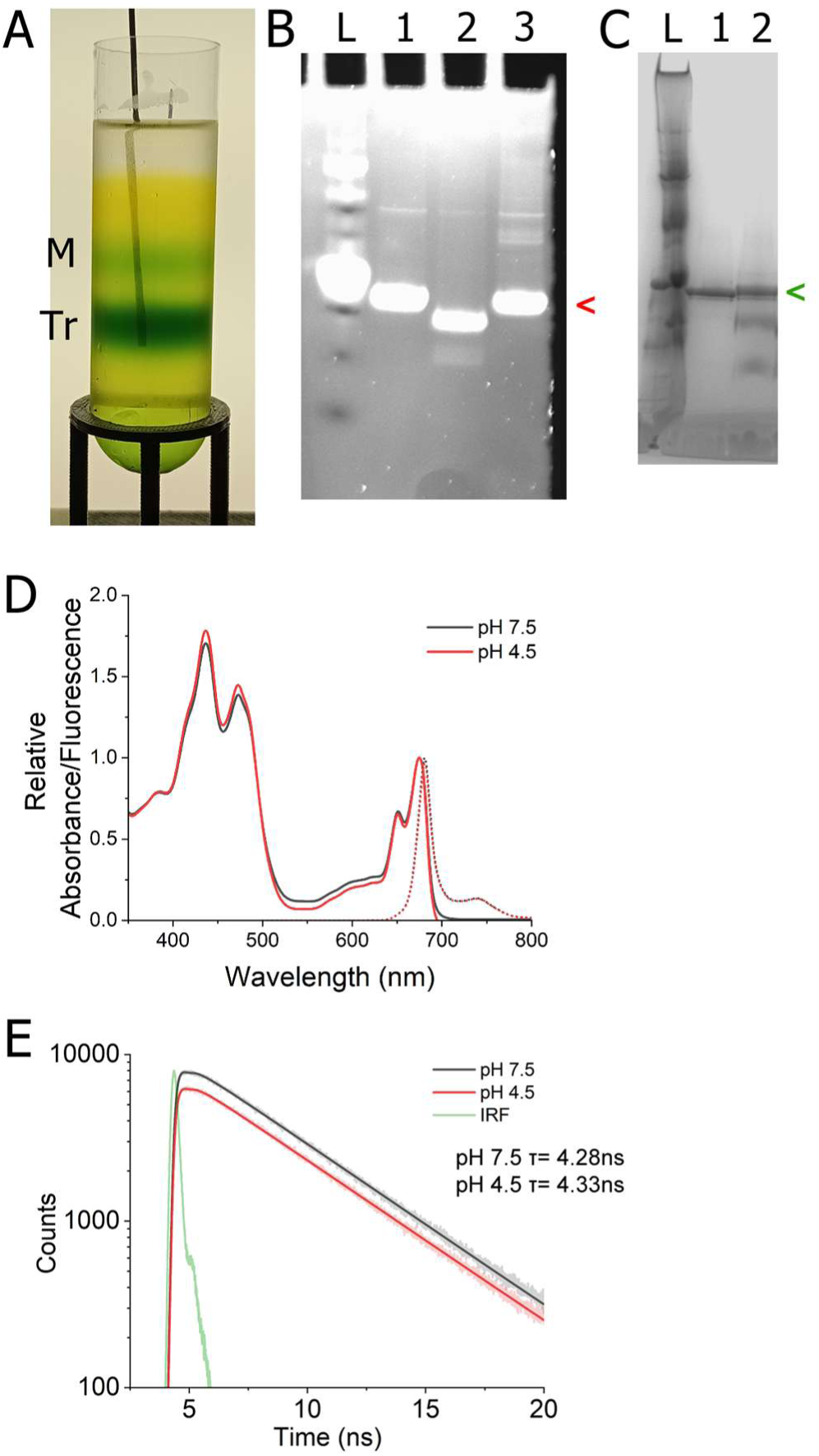
Spectroscopic analysis of LHCII at pH 7.5 and pH 4.5. (**A**) Example sucrose gradient (8-13.5% sucrose gradient, 0.03% α-DDM). The band representing LHCII trimers (Tr) was collected first, then the LHCII monomer band (M). (**B**) SDS-PAGE with SYPRO ruby stain. The gel lanes represent: Protein ladder (L), LHCII sample after SEC (1), Control sample of degraded LHCII stock after SEC (2), LHCII sample after sucrose gradient but before SEC (3). (**C**) Native-PAGE with Coomassie stain. The gel lanes represent: Ladder of crude thylakoid membranes (L), LHCII sample after SEC (1), LHCII sample after sucrose gradient but before SEC (2). (**D**) Absorbance (solid line) and emission spectra (dotted line, 436 nm excitation) of LHCII at pH 7.5 (black) and pH 4.5 (red). The absorbance spectrum was normalised to 1.0 at the 675 nm peak. The fluorescence emission spectrum represents the relative fluorescence corrected for different LHCII concentration by dividing by absorbance at 436 nm of the sample and displayed as a proportion of the emission of the pH 7.5 sample. See Fig. S3 for data from LHCII at all pH tested. (**E**) Fluorescence decay curve for LHCII at pH 7.5 (black). and pH 4.5 (red, translated by −2000 vertically, for visibility), after excitation with a 475 nm laser and collection at 680 nm. Fitted curves are displayed as solid lines, raw data as transparent lines. The instrument response function (IRF) is green. Graphs created with Origin 2025

### Single-particle cryoEM of LHCII – initial findings

After biochemical and optical characterization, the structure of LHCII was determined by single*-*particle cryoEM at both pH 7.5 and 4.5. Grids were prepared with carefully optimized blotting parameters and screened for the best conditions for ice quality and particle density. For LHCII at pH 7.5, a data set was collected of 5,374 micrographs at ×165,000 magnification and a total dose of 50.5 e^-^/A^2^. Full data collection statistics are available in Table S1. Data were processed using CryoSPARC software and two rounds of 2-D classification, with multiple rounds of non-uniform refinement, the initial particle stack of 2,786,556 was reduced to 118,642 for the final reconstruction which achieved 2.48 Å resolution, as determined by the gold standard FSC and a cFAR of 0.84 reflecting the good distribution of orientation (Fig. S1A, & and E) (Punjani et al., 2017). Fig. 2A and 2B show an initial overview of our raw electron density map and the atomic model (respectively) of the LHCII trimer solved at pH 7.5. Local resolution estimations indicate higher resolution of 2.2 Å around the important central region of each LHCII (Fig. 2A*, blue* colour in cutaway). A similar process was followed for the pH 4.5 data set, see Fig. S1B for details, but it became apparent there were two broadly different families of particles. The first family was very similar to that of the pH 7.5 structure with no apparent differences in either the secondary structure fold or the position of cofactors and made up approximately 25% of the data giving an overall resolution of 2.67 Å (Fig. S1B, D and F). The second family of structures showed clear changes consistent with destabilisation of the structure (approx. 75% of the data). Several maps could be produced, likely due to particles having a mix of one or two “destabilised” monomers within the trimer (no symmetry was applied during the structural refinement so each of the three monomers were independent). Ultimately, one map was chosen to represent the second family of LHCII at pH 4.5: this had one monomer matching the pH 7.5 map, one monomer showing the “destabilised” state and one monomer with a mixed density of the two with an overall resolution of 2.67 Å (Fig. S1B, D and F). The three different LHCII structures are described in more detail below.

**Fig. 2.**
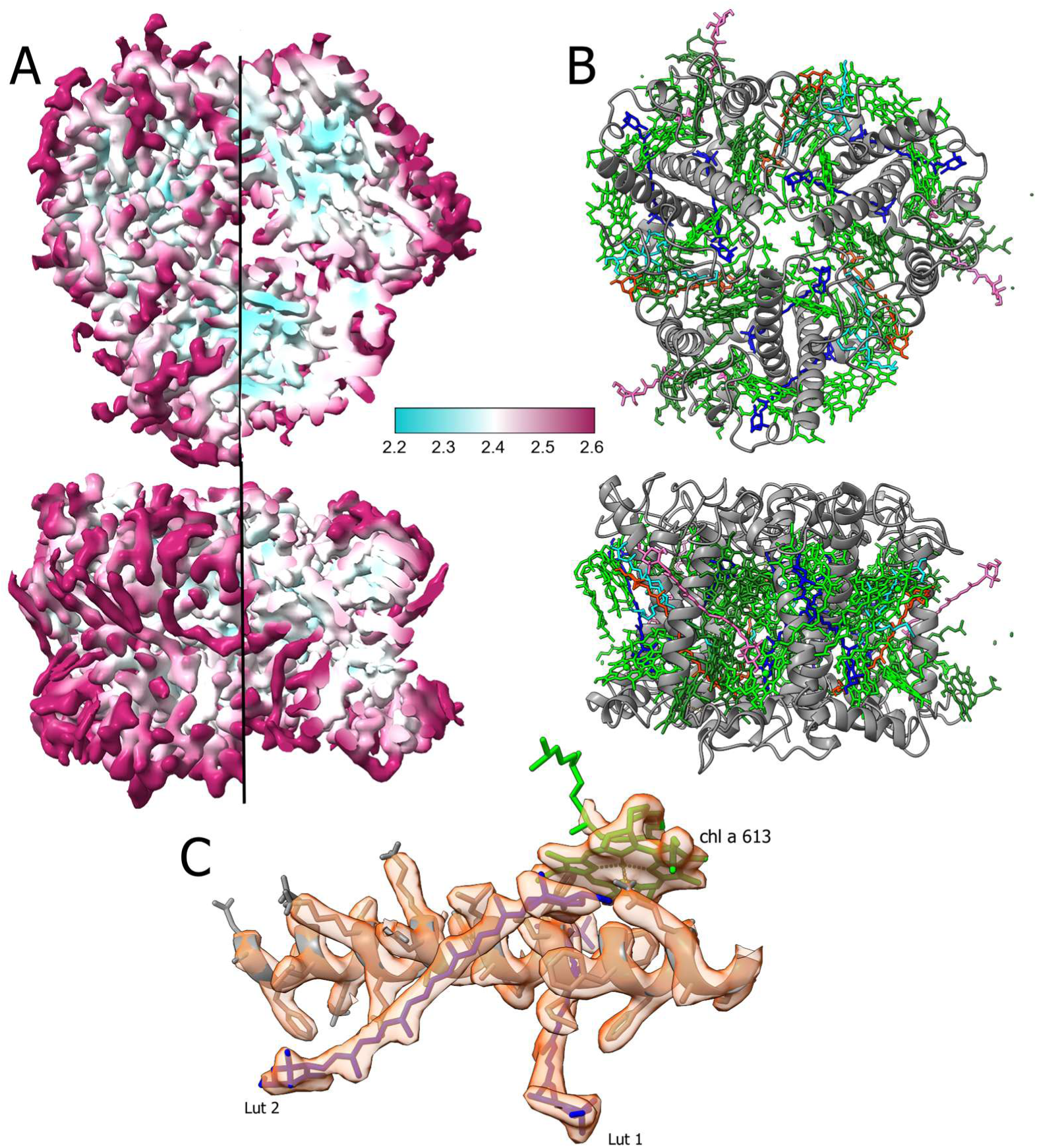
Overview of map and model for light harvesting LHCII in α-DDM at pH 7.5. (**A**) Electron density map of LHCII in α-DDM at pH 7.5, contour level 0.0768. Stromal side (top) and side view (bottom) coloured by local resolution in Å (see colour scale bar). The left side shows the electron density observable at the surface of the complex and the right side of each shows a slice into the density to show internal resolution of the complex. Overall resolution is 2.4 Å. (**B**) Atomic model of LHCII in α-DDM at pH 7.5 generated using the map in panel A. Polypeptide (grey); Chl *a* (light green); Chl *b* (dark green); Lutein (dark blue); Neoxanthin (pink); xanthophyll, modelled as Vio (orange), 1,2-Dipalmitoyl-Phosphatidyl-Glycerol (light blue). (**C**) An example region of the atomic model (wireframe) fitted into the electron density map (semi-transparent surface, orange) demonstrating the detail of the map and good fit of the model (contour level is 0.157)

### Structure of LHCII – in detail

The model of LHCII in α-DDM at pH 7.5 is broadly consistent with other published structures showing three transmembrane α-helices per monomer, two long transmembrane helices that cross the membrane at an angle, helix A and B, plus the third shorter, vertical helix E (Fig. 2B) (Kühlbrandt et al., 1994; Liu et al., 2004; Standfuss et al., 2005; Wang & Kühlbrandt, 1992). We follow the helix naming convention used for previous LHCII structures. The resolution achieved was such that we could accurately fit atomic models for most polypeptide side chains and all pigments (Fig. 2C). This contrasts with some previously published structures where the electron density was less defined so that the positioning of pigments was uncertain. The four Cars are clearly defined by the electron density and identifiable as follows: Lut in the L1 and L2 binding site, Neo in N1 binding site and Zea or Vio in V1 binding site (the Zea:Vio ratio was undetermined). The macrocycle ring of all eight chlorophyll *a* (Chl *a*) and six chlorophyll *b* (Chl *b*) molecules are very well-defined, and the majority of their phytol tails are clear with the exception of Chl *a* 614 and 604, plus Chl *b* 606 and 605, which have truncated tails in the map. Additional electron density is found within the map which would be consistent with either a bound lipid or detergent, however, as it is located on the edge of the structure and the identity is unclear, no molecule was fitted into this density when generating the atomic model (Fig. S4). Similar unidentified density was also found within the LHCII maps produced by Seki et al. (2024) suggesting this is a common feature of LHCII in α-DDM. Threefold rotational symmetry was observable in our maps without imposing any symmetry constraints and all three monomers have comparable density, combined with the 2-D classification classes (Fig. S1C and D), which is good evidence the map has been formed from intact trimers.

At pH 4.5, one of the two LHCII models obtained aligned with the pH 7.5 model described above without significant changes, as confirmed by a very low RMSD value of the polypeptide alpha carbons (RMSD-αC) of only 0.25 Å when comparing these two models (Fig. 3A). For the alternative LHCII pH 4.5 model, now referred to as the “destabilised pH 4.5 model”, one of the three monomers in the trimer appears to be destabilised, leading to a higher RMSD-αC of 0.62 Å for the trimer compared to the pH 7.5 structure, though this is mostly contributed to by the disordered C-terminus in the destabilised monomer (Fig. 3B, *red* region). Extensive processing of this region did not achieve a consensus map for the C-terminus beyond residue N223, suggesting it is highly dynamic. The density for the V1 car pigment that is clear in the pH 7.5 model is missing in the destabilized monomer of the pH 4.5 model (Fig. 3C). Furthermore, in the monomer lacking a V1 pigment, the Trp222 residue is rotated 180° and the Trp128 density disappeared in the neighbouring monomer, near the destabilised V1 site, suggesting disorder, though the α-helix it is part of remains well defined (Fig. 3D, see * and < annotations). The nearby Chl *a* 614 and the coordinating His212 in helix E of the destabilised monomer are also displaced (Fig. 3D) in the destabilised monomer at pH 4.5.

**Fig. 3.**
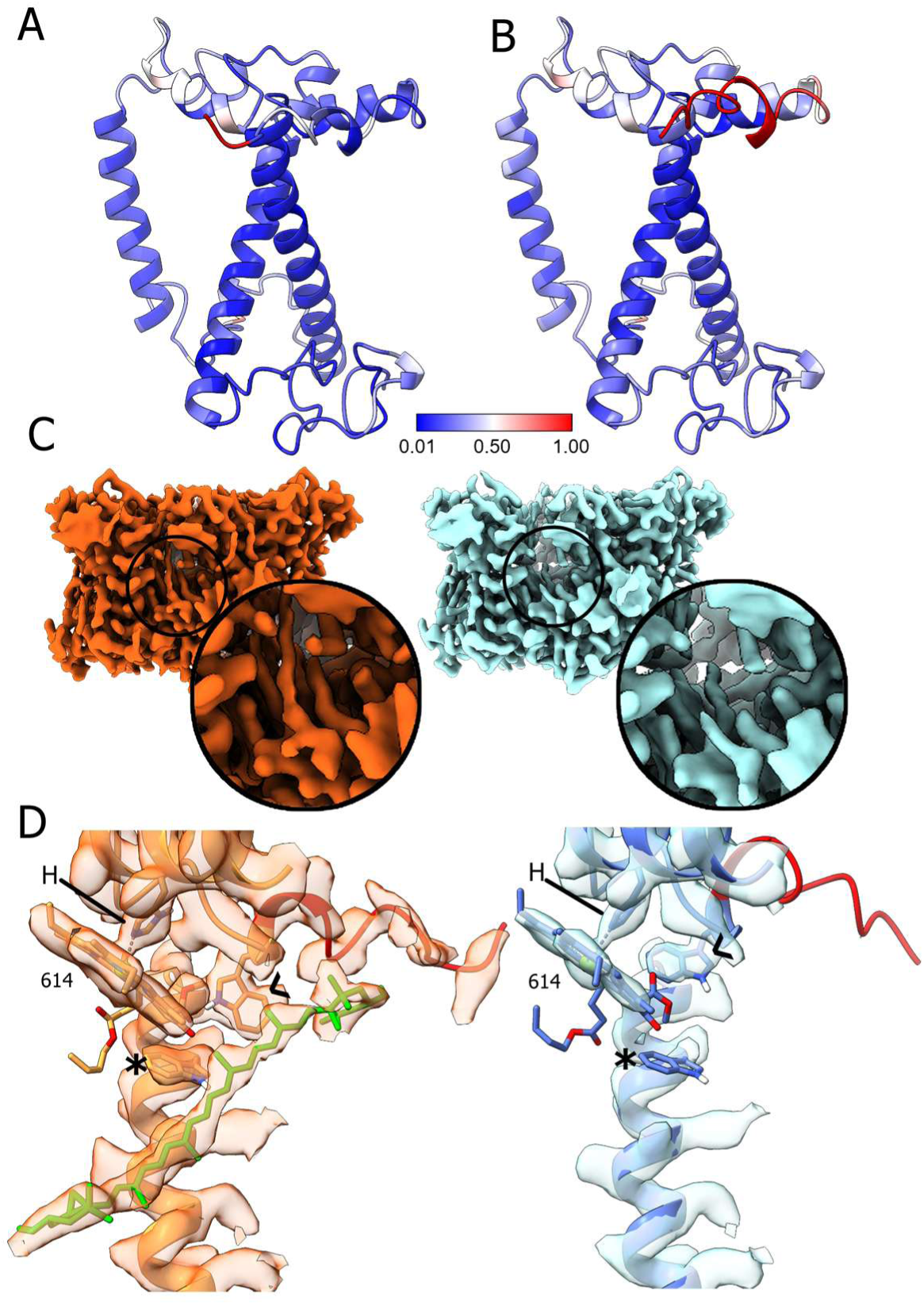
Comparison of models of LHCII-α-DDM at pH 7.5 and 4.5. (**A**) Model 1 (pH 4.5): one monomer of LHCII at pH 4.5, coloured by RMSD-αC in reference to the model of an LHCII monomer at pH 7.5. The overall RMSD-αC is 0.24 Å. Only the polypeptide is shown, for clarity. (**B**) Model 2 (pH 4.5): The destabilised monomer from the LHCII pH 4.5 destabilised model, coloured by RMSD-αC in reference to the pH 7.5 model. The overall RMSD-αC of the whole trimer is 0.29 Å, while the destabilised monomer displayed here has an RMSD-αC 1.07 Å but most of the difference is localised to the C-terminus (red) which lacks density in the map. (**C**) Electron density maps of LHCII at pH 7.5 (left, orange, contour: 0.0925) and the destabilized pH 4.5 map (right, blue, contour: 0.105). The V1 binding site is circled and shown enlarged in the inset. It is occupied by a xanthophyll pigment at pH 7.5 but the density is missing for the destabilised monomer. (**D**) Side-by-side comparison of the fitting of the model to the electron density around the V1 pigment binding site, for the “stabilized LHCII” versus “destabilised LHCII” structures. The LHCII at pH 7.5 model-in-map is shown in orange, left (contour 0.0848, 1.8 Å zone around visible atoms) compared to the “destabilised LHCII at pH 4.5” model-in-map shown in blue, right (0.109 contour, 1.8 Å zone around visible atoms). The V1 pigment (green) is missing from the destabilized pH 4.5 model, the Trp222 (<) is rotated 180°, the C-terminus (red) and Trp128 (*) of the neighbouring monomer is missing electron density. His212 (H) and *Chl a* 614 (614) are labelled

### Comparison of LHCII structural changes between pH 4.5 and 7.5

Previous literature on LHCII structure has suggested that a decrease in the distance between helix D and E drives a change in helix A/B crossing angle during the switch from light harvesting to energy quenching conformations (H. Li et al., 2020; Ruan et al., 2023). These reports suggested that a decreased helix crossing angle results in important pigment rearrangements, bringing Lut1 and the lowest energy Chl *a* trio closer, specifically, a decreased distance between Lut1 and Chl612 has been suggested as the ultimate trigger for quenching in the energy-dissipative state. ChimeraX was used to make distance measurements between specific pigments or residues, where pigments/residues-of-interest were chosen as they had been previously highlighted in other papers (analysis details provided in the Methods section and in Fig. S2). An overview of the pigment arrangement around the protein helices of LHCII is shown in Fig. 4A, together with schematics showing where the measurements were made for helix D-E separation (Fig. 4B), helix crossing angle (Fig. 4C) and Lut1 to Chl *a* 612 (Fig. 4D). The numerical data from the measurements are shown in Table 1. Each calculated distance (or angle) discussed below is the mean of the three monomers within each trimer, except for when referring to the “destabilised monomer” of the pH 4.5 model which is also noted individually. For LHCII at pH 7.5, helix D and E were found to have a separation of 5.3 Å and a helix crossing angle of 118*°* which is accompanied by a Lut1 to Chl *a* 612 separation of 5.5 Å (Table 1, bold). Both of the pH 4.5 models are very similar to the pH 7.5 model. The stable pH 4.5 model has a helix D to E separation of 5.4 Å and helix crossing angle of 118° with a Lut1 to Chl *a* 612 distance of 5.5 Å; only the helix D to E separation varies slightly compared to the pH 7.5 model. Meanwhile, the destabilised monomer of the pH 4.5 model has a helix crossing angle of 119°, marginally more obtuse, and a Lut1 to Chl *a* 612 separation of 5.3 Å (Table 1, bold). Since Chl *a* 610 and 613 are also situated closely to Lut1, their separation was also measured for all models. Distances between Lut1 and Chl *a* 610 and 613 were 5.6 Å and 4.8 Å, respectively, at pH 7.5 and both pH 4.5 models are similar (Table 1, bold). Overall, while there are some small changes between models, this variation tends to be no larger than the variation between monomers of the same model suggesting no significant difference. Each monomer within the trimer was modelled independently from the raw EM data and the low standard deviations of these measurements show that there is little variation between them.

**Fig. 4.**
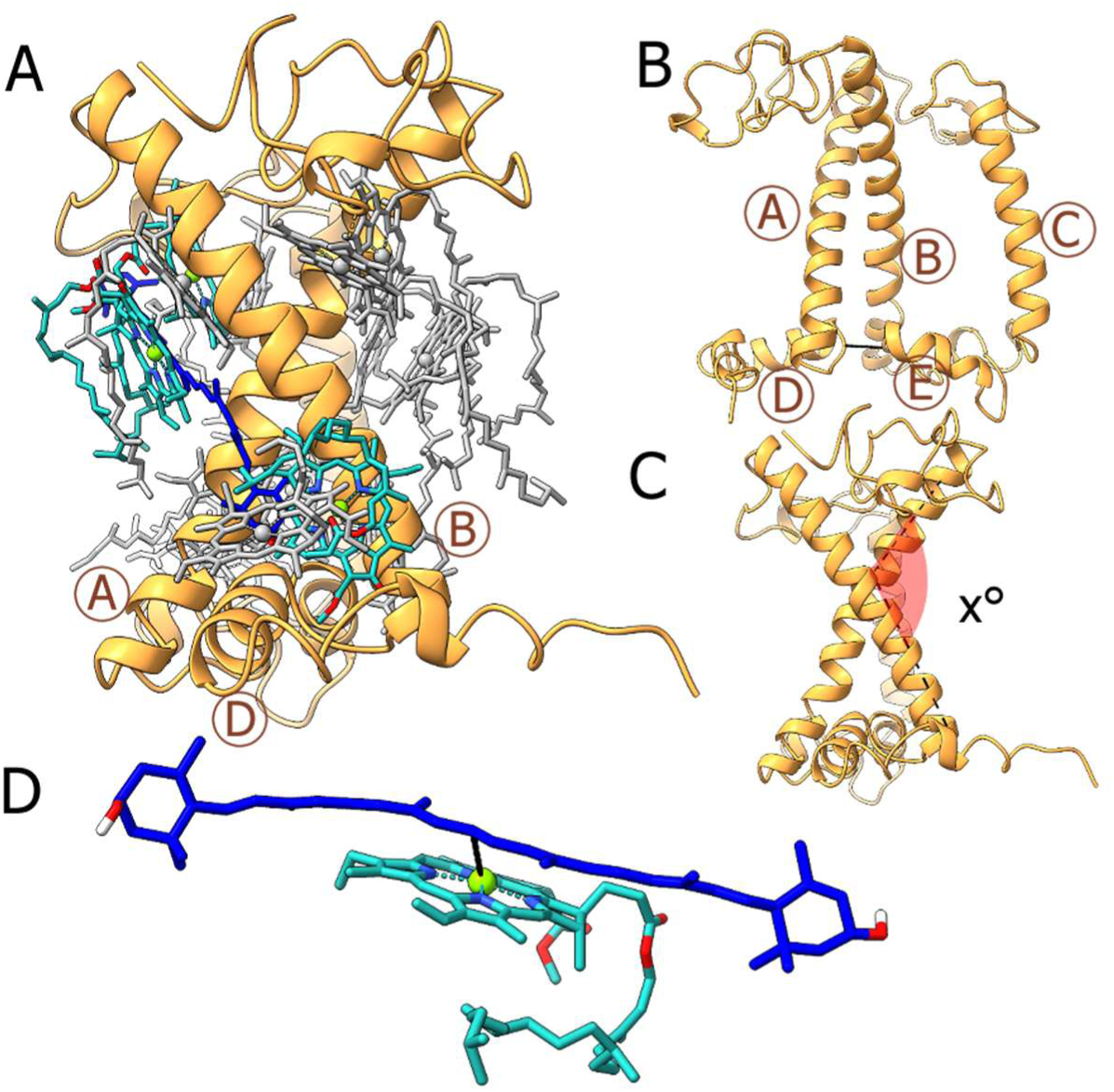
Views of the atomic model of LHCII at pH 7.5 showing how measurements of distances and angles of interest were made. (**A**) The arrangement of Chl around helix A and B. The lowest energy trio are coloured in blue, the other Chl are grey. (**B**) Arrangement of helices A-E of one LHCII monomer. The dotted line shows the separation distance measured between E96 and V204 of helix D and E. (**C**) View of helix A and B where the crossing angle, x°, is indicted by a dotted line and red shading. It is defined by the angle between alpha carbons of F58, C69, R185 and V200. (**D**) View of the relative positions of Lut1 and Chl 612, highlighting the separation distance that we define by the C15 of Lut to the Mg of Chl 612. Helix labels are circled and in brown

**Table 1.**
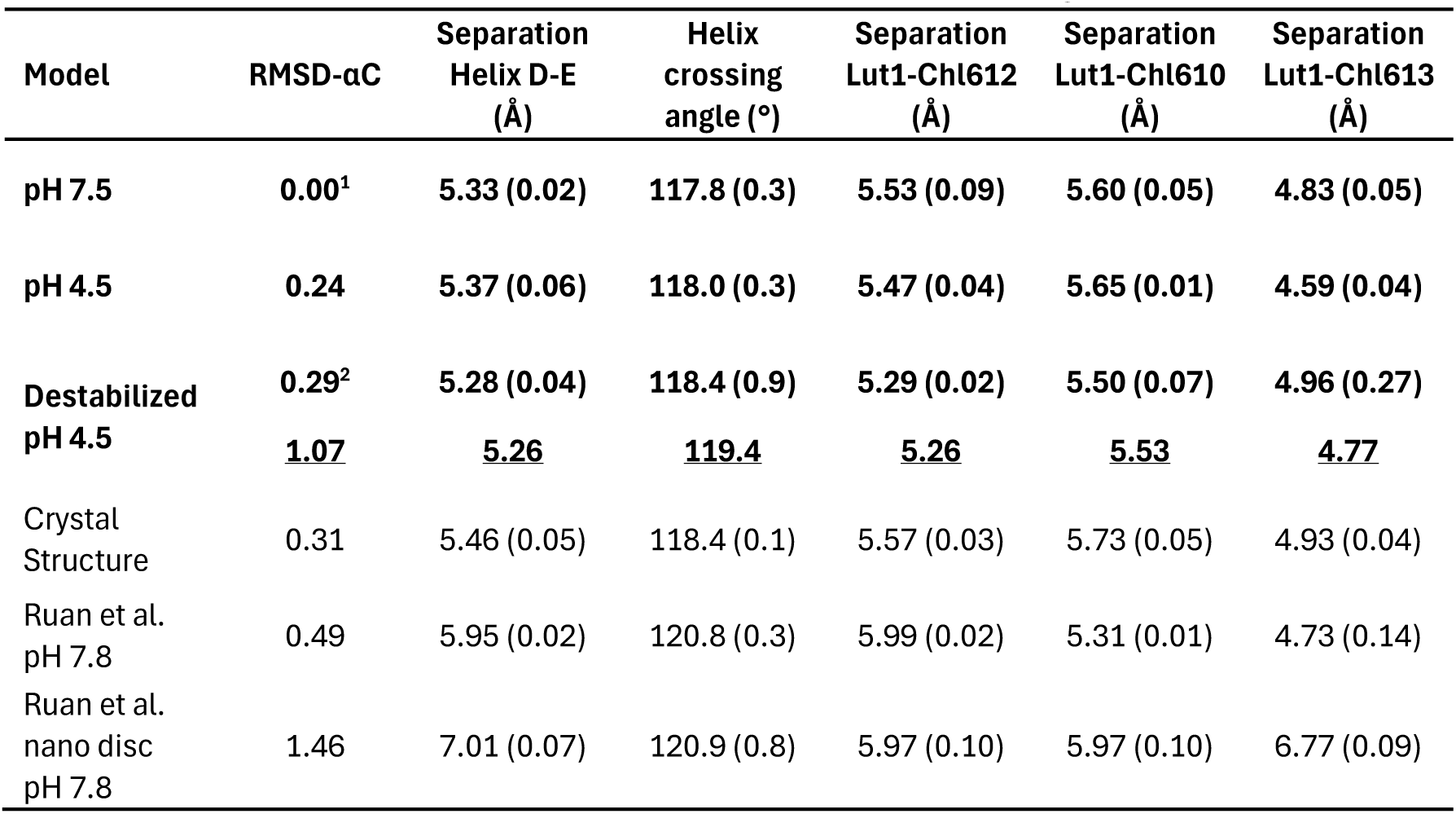
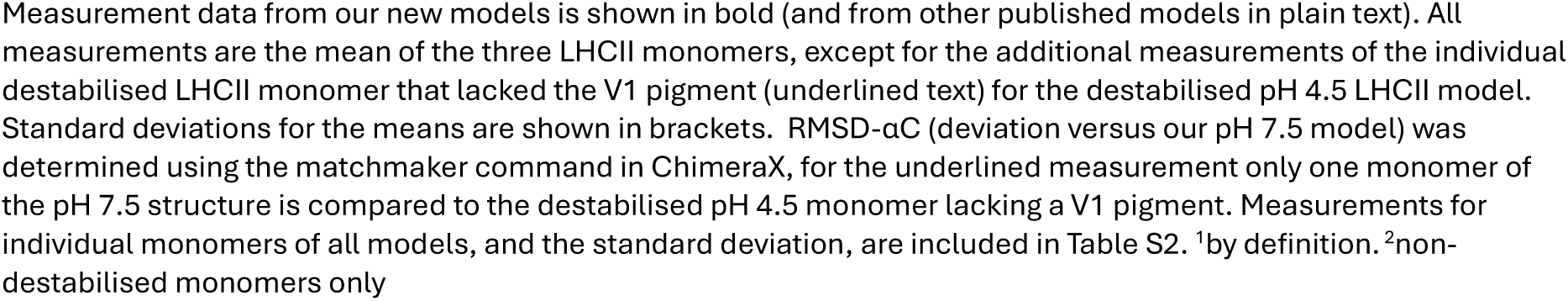
Measurements of various atomic models of LHCII to compare inter-pigment and intra-protein architecture.

| Model | RMSD-αC | Separation<br>Helix D-E<br>(Å) | Helix<br>crossing<br>angle ($^\circ$) | Separation<br>Lut1-Chl612<br>(Å) | Separation<br>Lut1-Chl610<br>(Å) | Separation<br>Lut1-Chl613<br>(Å) |
| --- | --- | --- | --- | --- | --- | --- |
| pH 7.5 | 0.00<sup>1</sup> | 5.33 (0.02) | 117.8 (0.3) | 5.53 (0.09) | 5.60 (0.05) | 4.83 (0.05) |
| pH 4.5 | 0.24 | 5.37 (0.06) | 118.0 (0.3) | 5.47 (0.04) | 5.65 (0.01) | 4.59 (0.04) |
| Destabilized<br>pH 4.5 | 0.29<sup>2</sup> | 5.28 (0.04) | 118.4 (0.9) | 5.29 (0.02) | 5.50 (0.07) | 4.96 (0.27) |
|  | <u>1.07</u> | <u>5.26</u> | <u>119.4</u> | <u>5.26</u> | <u>5.53</u> | <u>4.77</u> |
| Crystal<br>Structure | 0.31 | 5.46 (0.05) | 118.4 (0.1) | 5.57 (0.03) | 5.73 (0.05) | 4.93 (0.04) |
| Ruan et al.<br>pH 7.8 | 0.49 | 5.95 (0.02) | 120.8 (0.3) | 5.99 (0.02) | 5.31 (0.01) | 4.73 (0.14) |
| Ruan et al.<br>nano disc<br>pH 7.8 | 1.46 | 7.01 (0.07) | 120.9 (0.8) | 5.97 (0.10) | 5.97 (0.10) | 6.77 (0.09) |

**Fig. 4** Views of the atomic model of LHCII at pH 7.5 showing how measurements of distances and angles of interest were made. (**A**) The arrangement of Chl around helix A and B. The lowest energy trio are coloured in blue, the other Chl are grey. (**B**) Arrangement of helices A-E of one LHCII monomer. The dotted line shows the separation distance measured between E96 and V204 of helix D and E. (**C**) View of helix A and B where the crossing angle, x°, is indicted by a dotted line and red shading. It is defined by the angle between alpha carbons of F58, C69, R185 and V200. (**D**) View of the relative positions of Lut1 and Chl 612, highlighting the separation distance that we define by the C15 of Lut to the Mg of Chl 612. Helix labels are circled and in brown

### New insights into pigment conformations

The conformation of the Lut pigments within LHCII are key interests as they likely play an important role in switching between the light harvesting and the quenching state (Accomasso et al., 2024; Holleboom & Walla, 2014; Pedraza-González et al., 2024). Previous cryoEM models of LHCII had diffuse electron density around these pigments so that conformation of the Car in the model was unclear (Fig. S5 and Fig. S6) (Ruan et al., 2023). In contrast, our new models have accurately positioned Lut (and other Cars) due to tight electron densities around all regions of the pigments, allowing unambiguous placement of these structures into the model. The resolution of the map of LHCII at pH 7.5 near to the L1 and L2 Lut binding sites are ∼2.2 Å (Fig. 2A), allowing the unambiguous positioning and orientation of Lut within the structural model fitted to this data (Fig. 2C). Molecular simulations predict a different conformation for each Lut, and these different conformations are thought to impart access to different energy levels according to quantum chemical calculations (Accomasso et al., 2024; Fox et al., 2015; J. Li et al., 2023; Macernis et al., 2012). Our high-resolution map clearly defines the Lut conformations and reveals twists in the backbone leading to relatively different positions of the Car chain coordinates. A visual comparison of the molecular geometry of these two Cars was carried out by overlaying Lut1 and Lut2 of the pH 7.5 model so that the middle of the chains were aligned (Fig. 5A, region III, grey): the conformation within this region has little variation between Lut1 and Lut2 (within one model) so this region was taken as a reference point. Torsion measurements were also made using ChimeraX to quantify the rotation of region II and IV (Fig. 5A, red regions) and the rotation of each end ring in regions I and V (Fig. 5A, boxed regions). For an illustration of the carbon numbering system used and all measurements taken for the lutein molecules: see Fig. S5. At pH 7.5, focussing on the lumenal side of the Car molecule, region II, Lut1 is found to be comparatively more twisted than region II of Lut2, which is clear from a visual comparison of the maps (see Fig. 5B, * annotations). Quantification of these changes via measurements show that Lut1 twists 18° while Lut2 twists only 6° (Table 2, in bold). Meanwhile, region IV, the stromal side of the Car, is more rotated in Lut2, at 25° compared to 17° in Lut1 though neither Lut is straight in this region (Fig. 5C and Table 2, in bold). The trend for the Lut2 conformation is consistent between all three models, with significant twisting only occurring in region IV, although both pH 4.5 models show more rotation of region IV than the pH 7.5 model, while region II is less twisted in the pH 4.5 model (3°) and more twisted in the destabilised pH 4.5 monomer model (9°) compared to the pH 7.5 model (6°). Meanwhile, the Lut1 conformation is disturbed in the destabilised pH 4.5 model with more variation between monomers, shown by the higher standard deviation, in region IV (Table 2, bold brackets). In the destabilised monomer, the Lut1 region IV rotation is increased to 20° whilst the average of all three trimers was relatively lower (straighter) on both the lumenal and stromal sides overall (Table 2, bold). One may have expected that within the destabilized pH 4.5 LHCII trimer, the destabilised monomer with the missing V1 pigment would show the greatest change in pigment conformation as compared to the other two models of stable LHCII. Interestingly, this was not the case for all measurements, which could reflect a more far-reaching effect of an empty V1 binding site in one LHCII monomer changing the structure in the other two monomers within the LHCII trimer. In summary, these detailed measurements of the Lut chain twist and headgroup rotation revealed subtle differences between Lut1 and Lut2 within one LHCII and elucidated the similarities and differences in Lut conformations caused by a pH change.

**Fig. 5.**
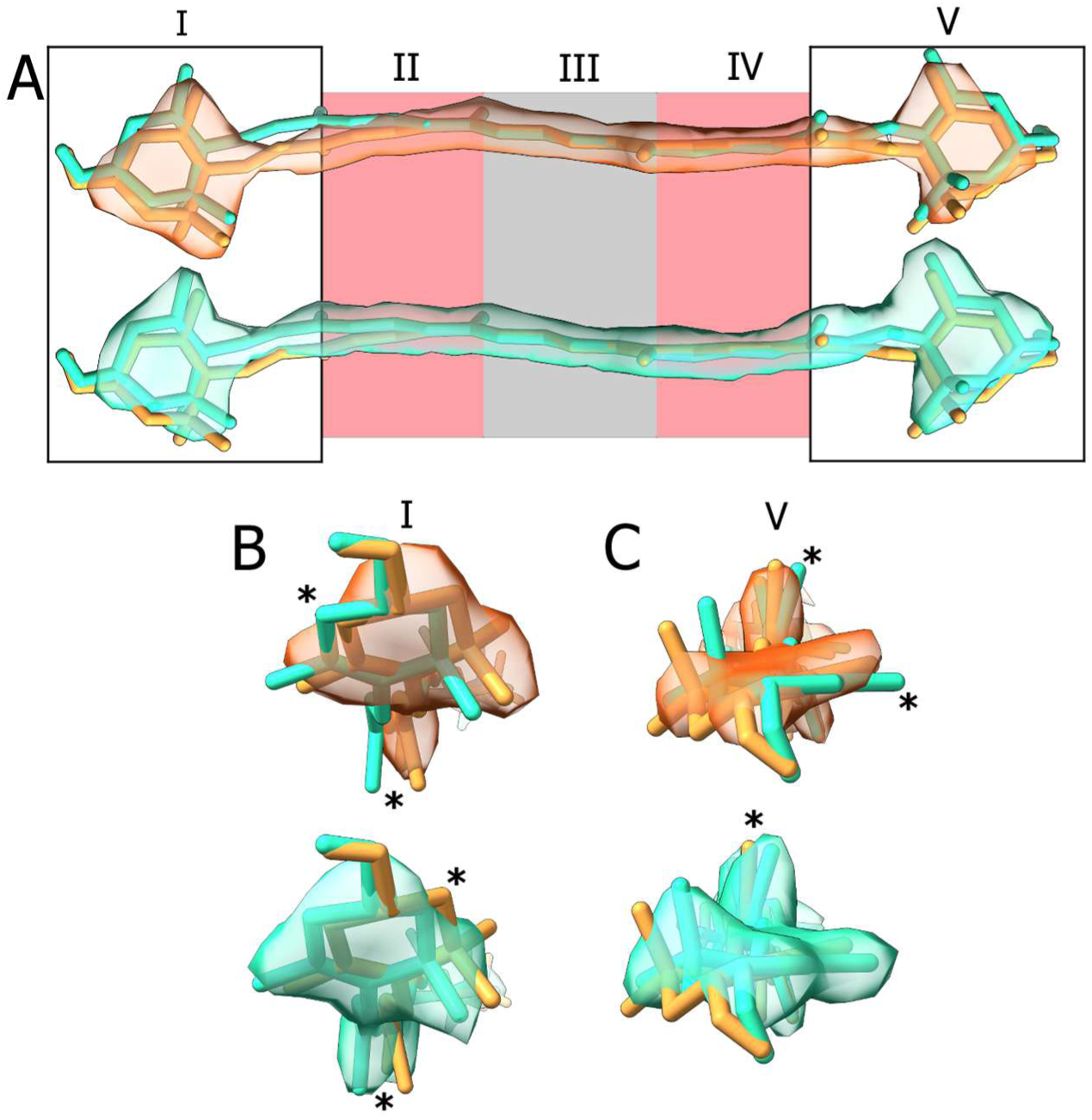
The two luteins have different conformations which are clearly defined by the map. (**A**) Lut1 (orange) and Lut2 (blue) of the pH 7.5 model are overlaid so region III (grey box) aligns - the conformation in this region is near identical for both Luts (side-on). Region I is the lumenal side, region V is the stromal side. Regions II and IV (red) were measured for rotation. (**B**) View of region I, along the length of the molecules (end-on). Lut1 has distinctly more of a twist compared to Lut2, resulting in clear regions of Lut1 that will not fit in the Lut2 map and visa-versa (noted by *) (**C**) View of region V, along the length of the molecules (end-on). Lut2 has more of a twist on stromal side than Lut1, though more subtle compared to the lumen side, leading to regions of the atomic structures that will not fit in the other map (noted by *). For each panel, top: both Lut molecular models in the Lut1 electron density map (orange), bottom: molecular models of both Luts with only Lut2 electron density map (blue). Lut1 in the context of its binding site can be seen in Fig. S7

**Table 2.** Quantifying lutein conformations.

| Model | Lut1 twist<br>region II<br>(°) | Lut1 twist<br>region IV<br>(°) | Lut1 ring<br>rotation<br>region I (°) | Lut1 ring<br>rotation<br>region V (°) | Lut2 twist<br>region II<br>(°) | Lut2 twist<br>region IV<br>(°) | Lut2 ring<br>rotation<br>region I (°) | Lut2 ring<br>rotation<br>region V (°) |
| --- | --- | --- | --- | --- | --- | --- | --- | --- |
| **pH 7.5** | **17.9 (1.4)** | **16.5 (1.4)** | **64.1 (2.4)** | **117.2 (4.8)** | **5.5 (3.4)** | **24.6 (2.7)** | **75.3 (1.2)** | **139.2 (1.1)** |
| **pH 4.5** | **16.2 (3.1)** | **17.6 (1.8)** | **67.3 (3.2)** | **124.9 (1.3)** | **3.0 (1.4)** | **28.6 (5.1)** | **70.5 (1.0)** | **136.4 (0.4)** |
| **Destabilized**<br>**pH 4.5** | **11.2 (3.0)** | **11.8 (5.9)** | **98.7 (0.6)** | **125.3 (1.9)** | **7.6 (1.6)** | **27.4 (5.2)** | **64.5 (1.4)** | **138.3 (2.6)** |
|  | <u>13.3</u> | <u>20.1</u> | <u>98.7</u> | <u>127.5</u> | <u>9.0</u> | <u>27.4</u> | <u>66.43</u> | <u>139.1</u> |
| crystal<br>structure | 8.9 (0.2) | 3.9 (0.7) | 77.0 (0.3) | 121.2 (1.2) | 1.6 (1.4) | 10.8 (0.4) | 80.2 (1.1) | 127.6 (1.2) |

## Discussion

Here we report new structural models of the light harvesting state of LHCII, determined by single-particle cryoEM, whose map quality allows significant new molecular details to be elucidated (Fig. 2 and Fig. 3). The high-resolution structures of LHCII at both pH 7.5 and 4.5 allowed us to unambiguously fit all pigments of LHCII into three electron density maps, one produced at pH 7.5 and two others at pH 4.5. Of the two LHCII models at pH 4.5, one matches the pH 7.5 model and the other is representative of a destabilised state with one monomer containing a disordered C-terminus, missing the V1 Car and with a shift in helix E, induced by low pH (Fig. 3). Based on particle numbers, the majority of LHCII trimer particles in this low-pH sample have at least one destabilised monomer, yet, the bulk fluorescence lifetime and emission were not reduced (Fig. 1E). There are subtle changes of the pigment arrangement, most notably, loss of the V1 (Zea or Vio) pigment, changes to Lut1 conformation and somewhat reduced separation between Lut1 and nearby Chl *a*. The similarity in the fluorescence lifetimes between the pH 7.5 and pH 4.5 samples, despite these structural changes, highlights that they are insufficient to induce the quenched state. Later in this Discussion, we compare our new LHCII models to previously solved structures to identify new insights provided by our higher resolution.

Previous work suggested that the crystal (X-ray) models of LHCII must represent the energy-dissipative state due to their short fluorescence lifetime (Pascal et al., 2005). Meanwhile, single-particle cryoEM has been used to resolve models from samples with both long, light-harvesting-like fluorescence lifetimes and somewhat shorter, quenching-like lifetimes (Ruan et al., 2023). In the current report, all of our samples possess >4ns lifetimes, characteristic of the light harvesting state. Therefore, comparison of the structural differences between our new structural models and previous models should be highly informative. In the case of Ruan et al, LHCII structures were resolved in both detergent micelles (β-DDM) and lipid nanodiscs at pH 7.5 and pH 5.4 via cryoEM, achieving models of the “unprotonated” and “protonated” structures where subtle conformational differences between the two were suggested to represent light harvesting LHCII and the quenched LHCII, respectively (Ruan et al., 2023). However, while the overall reported resolution was similar to the new structures presented here, at 2.5-2.8 Å, there was a notable decrease in the quality of the electron density map towards the centre of each monomer for the Ruan et al. maps, especially around the Cars, where small changes in separation distance or conformation may have a vital impact on the energy transfer within the trimer (Liguori et al., 2015). A side-by-side comparison of the electron density for regions-of-interest around the most important pigments are shown in Fig. S5 for neutral pH maps and Fig. S6 for low pH maps where it is clear that the electron density is relatively tight (i.e., of similar certainty) for the current LHCII structure and the crystal structure (Lui et al. 2012) but the electron density is more diffuse for the Ruan et al. (2023) structures. Each of these structures were excellent advances in their own time, with particular novelty in the use lipid nanodiscs (Ruan et al., 2023), but it is fair to revisit them and compare against our higher resolution data. In other work, Seki et al. (2024) published two single particle cryoEM maps of natural LHCII trimers extracted in α-DDM, one with C3 symmetry imposed (2.17 Å) and the other without imposing symmetry (2.32 Å), including a model for the C3 map. The map quality is high throughout (similar quality to our data), however, no fluorescence lifetime was reported. While it is likely to be light harvesting, as the LHCII was isolated in α-DDM at pH 7.5 as ours was, it cannot be confirmed. Multiple X-ray crystallography structures of LHCII trimers have also been published, all with highly similar structures and short fluorescence lifetimes indicative of energy quenching state (Dockter et al., 2012; Pascal et al., 2005). In our later comparisons, we will focus on the model by Liu et al. as it has the highest resolution and most complete map, and it will be referred to as “the crystal structure” hereafter (Liu et al., 2004).

### Comparison of the overall polypeptide structure found in LHCII crystals versus isolated LHCII

LHCII crystals were reported to have short fluorescence lifetimes of <0.5 ns, indicative of energy quenching. Our new model, based on LHCII that is isolated as single proteins within detergent micelles and has long lifetimes of >4 ns, provides an opportunity to explore the structural reasons for energy quenching in LHCII. It should be noted that LHCII within crystals are in relatively close contact with one another, and LHCII-LHCII interactions could potentially lead to other pathways of energy dissipation (e.g., via singlet-triplet and triplet-triplet annihilation) that mimic the decreased fluorescence lifetimes that are expected for an energy-quenching conformation of LHCII, without changing the internal conformation of individual LHCII complexes (Barros et al., 2009; Conradie et al., 2026; Manna et al., 2023; Satpathi et al., 2025). Thus, questions remain around the true quenched state of an individual LHCII. An aggregation-based quenching may be responsible for the reduced fluorescence lifetime of crystalline LHCII, but single-particle fluorescence studies have shown that isolated LHCII can also fluctuate between a normal and a reduced fluorescence lifetime (Chmeliov et al., 2019; Gao et al., 2025; Schlau-Cohen et al., 2015). This implies the existence of a mechanism of LHCII quenching without aggregation, potentially involving a conformational change, and it is possible there are separate mechanisms which occur in vitro but may or may not occur in the native thylakoid membranes.

Fig. 6 and Tables 1-2 provide visualizations and the results of measurements that compare our new LHCII models to pre-existing LHCII models. In our measurements, we find that the new LHCII structures solved by single-particle cryoEM align closely with the previous crystal structure, in contrast to the single-particle cryoEM models of Ruan et al. (2023), either in β-DDM or in lipid nanodiscs, which are substantially different. Moreover, our model is also in much better agreement with the single-particle cryoEM model of LHCII solved by Seki et al. (2024), also in α-DDM. Comparing our LHCII (pH 7.5) model to others gives polypeptide backbone deviations (RMSD-αC) of 0.36 Å, 0.34 Å and a range 0.53-2.27 Å, for the crystal structure, Seki et al. structure and Ruan et al. structures, respectively (see Fig. 6A), in other words, they are similar, similar and variable, respectively. Furthermore, the variation between structures tends to be greater towards the exterior of the model, where the protein is naturally more mobile (Dockter et al., 2012), rather than the regions closest to expected site of quenching (Fig. 6A). Overall, our new LHCII models are more similar to the crystal structures than to the LHCII models reported by Ruan et al. (2023) (Table 1) This supports the conclusion that our new models and the crystal structure share very similar polypeptide architecture, despite the differing fluorescence lifetimes, raising questions about how the difference in energetics may come about. Later in the Discussion, we delve into the similarities and differences in specific regions of the LHCII structure that could have an impact on NPQ/qE.

**Fig. 6.**
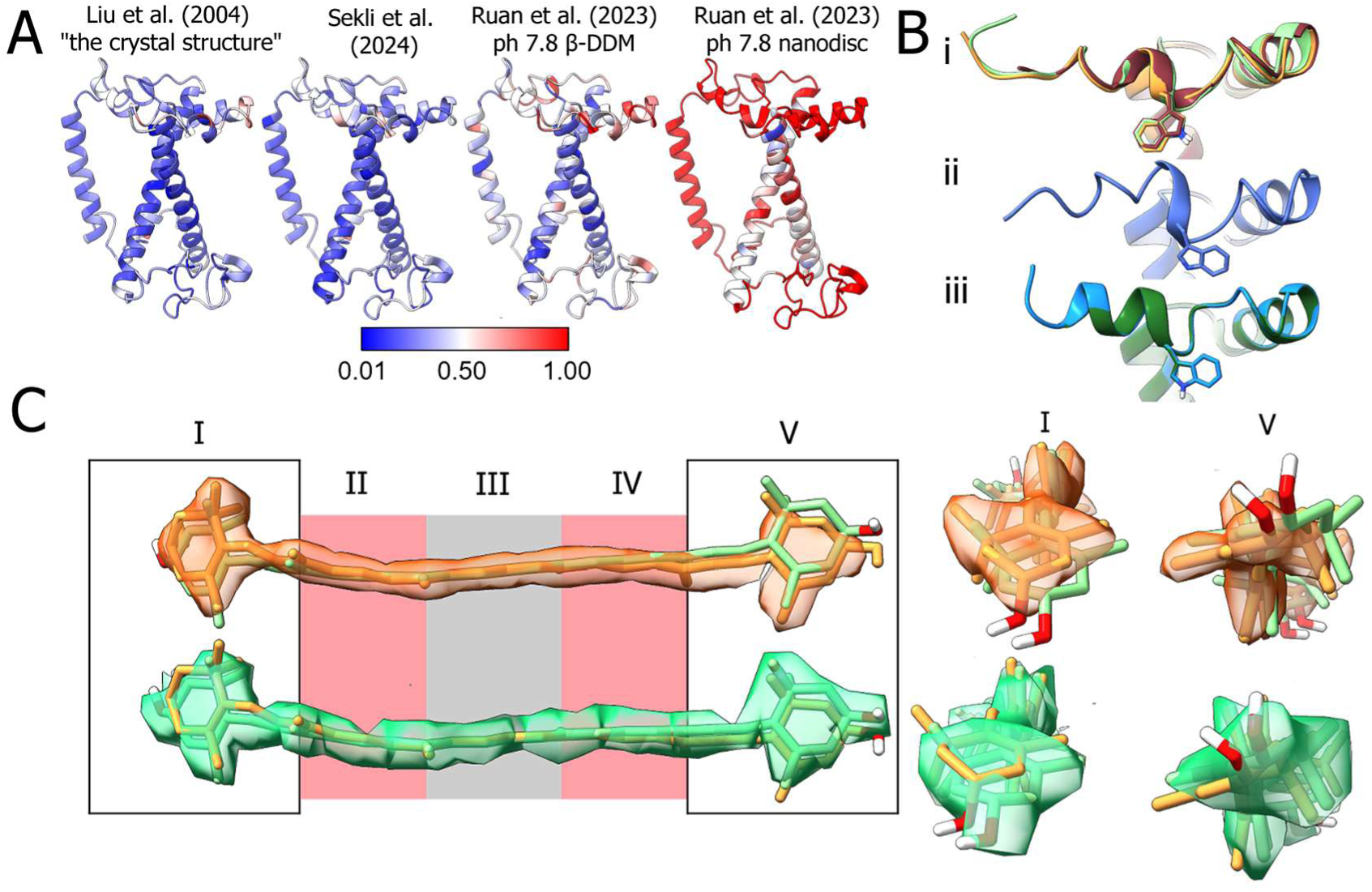
Comparison of published LHCII structures. (**A**) Atomic models of LHCII coloured by RMSD-αC in comparison with our pH 7.5 model. From left to right: the Liu et al. crystal structure of LHCII, Seki et al. natural LHCII, Ruan et al. pH 7.8 LHCII in β-DDM, Ruan pH 7.8 LHCII in nanodiscs. (**B**) Comparison of atomic models of the C-terminus of LHCII, focussing on the orientation of Trp222 with three groups of structures overlaid, for clarity, in each. Group (i) with Trp222 orientated towards the ordered C-terminus where our pH 7.5 LHCII model is in orange, the Seki et al. model in brown, the crystal structure model in pale green. Group (ii) with Trp222 orientated away from a destabilized C-terminus showing our destabilised LHCII monomer at pH 4.5 in blue. Group (iii) Trp222 orientated away from the α-helical C-terminus showing the Ruan et al. LHCII model at pH 5.4 in β-DDM detergent in blue and in nanodiscs in green. The Trp222 side chain is rotated 180° in groups ii and iii compared to group i. (**C**) Overlay of Lut1 from our pH 7.5 model (orange map and model, contour 0.193) and the crystal structure (green map and model, contour 0.291), aligned in region III to show the different conformations. Views of the Lut molecule are shown either: side-on showing the length of the backbone (left), or end-on from region I (middle), or end-on from region V (right). Comparisons to crystal structures of LHCII from Standfuss et al. and Wang et al. were also made and, as anticipated, generally align with the Liu et al. crystal structure (Standfuss et al., 2005; Wang & Kühlbrandt, 1992). A complete table of measurements is provided in Table S2 and the Lut1s in the context of their binding sites can be seen in Fig. S7

### The structure of the C-terminus of LHCII

At pH 4.5, we observed that in some monomers of the trimer, the Trp222 is rotated and the remaining C-terminus became disordered, the V1 Car is also lost in these monomers (Fig. 3D). It has been previously observed in LHCII, that disorder in this region leads to loss of the V1 Car, highlighting the importance of the C-terminus in binding the V1 Car (Z. Li et al., 2023; Seki et al., 2024). In the thylakoid, when pH drops during exposure to high light intensity, the pigment at this binding site is converted from Vio to Zea by the enzyme violaxanthin de-epoxidase and appears to allow LHCII to respond to physiologically-relevant lowered pH (Goss et al., 2017). It is currently unclear how the enzyme can access the pigment in LHCII as it is not exposed to the solvent. It is possible that the pH-dependant C-terminus disorder that we are highlighting in our LHCII models would allow an enzyme access to the pigment, however, we do not have enough evidence to be clear whether this event is a physiological response to pH or an artifact of the detergent/purification processes. Comparing our new LHCII models to previously published structures, we find that our pH 7.5 model and “ordered pH 4.5 model” align with other neutral pH structures in this region: Trp222 consistently orientates in the same way and the remaining C-terminus is an ordered coil (Fig. 6B, group I). Once the pH drops, in at least some monomers, Trp222 rotates 180°, shown in both our “destabilized pH 4.5 model” (Fig. 6B, group ii) and low pH models from Ruan et al. (Fig. 6B, group iii). However, in contrast to our findings, Ruan et al. report the formation of an additional α-helix at the C-terminus with convincing density (Fig. 6B, group iii), also supported by molecular simulations and 2-D infrared spectroscopy of LHCII in α-DDM at pH 4, which suggest an increase in α-helical content at low pH (Do et al., 2026; H. Li et al., 2020; Ruan et al., 2023). The physiological relevance of this reorientation of the C-terminal Trp remains unclear.

### Analysis of the Lut1-Chl separation distances between different LHCII models and the effect on quenching

Previous studies have suggested that an important structural rearrangement in the change of LHCII from a light harvesting state to a quenched state involves: (i) the inward motion of helix D and E, (ii) the angle of the crossing helices A and B becoming more acute, (iii) a decreased separation between Lut1 and Chl *a* 612. Ultimately, these changes have been suggested to cause increased energy dissipation by Lut1 (Daskalakis et al., 2019; Ruan et al., 2023; Wilson et al., 2024; Yan et al., 2007). Measurements of the helix crossing angles, helix separations and the related Chl-Lut inter-pigment distances were made to compare our new LHCII structures to previously published ones (see Table 1 and Fig. 4). Our helix crossing angles (∼118°) align with the crystal structure (∼118°) compared to Ruan’s proposed light harvesting structure (∼121°), our helix D-E separation distances (5.3-5.4 Å) are similar to the crystal structure (∼5.5 Å) as compared to longer distances in the Ruan light harvesting structure (∼7.0 Å), and our Lut1 to Chl 612 separation is also similar to the crystal structure (5.3-5.5 Å compared to 5.6 Å). These measurements are reproducible and have relatively low standard deviations. The lack of significant differences between our new LHCII structures and the crystal structure, would imply, if the earlier hypothesis is correct, that they should have a similar energetic state. Therefore, there is an interesting discrepancy in the energetic states expected for LHCII, between these different models, which has important consequence and a few possible explanations. One possibility is that our new LHCII model and the crystal structure model both represent an energy-dissipative state and the Ruan model represent the light harvesting state. This seems unlikely because it is improbable that our LHCII sample would become quenched during preparation for EM grids but the Ruan LHCII sample did not (sample preparation for cryoEM is standardized). A second possibility is that our new LHCII model and the crystal structure model both actually represent a light harvesting state and the Ruan model is anomalous due to the limitations of their more diffuse electron density map. This could be possible if the reduced fluorescence lifetime observed for the crystal structure of LHCII is actually caused by artefactual exciton annihilation effects rather than a (true) physiological dissipative state. Recent studies have shown that singlet-triplet annihilation effects can greatly reduce the fluorescence lifetime of LH complexes simply due to the connectivity of neighbouring proteins and may not necessarily represent an internal structural change to the LHCII protein (Conradie et al., 2026; Manna et al., 2023; Satpathi et al., 2025). A third possibility is that our new LHCII model and the crystal structure do represent the light harvesting and quenched states, respectively, as expected from their fluorescence lifetimes, which implies that the different energetic states do not require significant differences in polypeptide structure. Either the differences in 3-D protein structure are (i) very subtle, (ii) do not exist, or (iii) cannot be captured by current structural techniques. Perhaps, subtle differences in the pigment configurations could occur and be sufficient to drive a dissipative state of LHCII, despite similar polypeptide architectures. The next section of the discussion explores this possibility.

### Differences in lutein conformation between different LHCII models and the effect on quenching

Having considered how the polypeptide structure may induce changes in the pigment-pigment distance we now consider the Lut pigments in greater depth. Carotenoids, including Lut, have complex photophysics, making them suitable for both light harvesting and energy quenching. The Lut1 within LHCII is commonly regarded as the site of final quenching (Ilioaia et al., 2011; Son et al., 2019; Yan et al., 2007). Ultrafast spectroscopy data and theoretical modelling have led to debate over of Lut energetics within LHCII. It is generally accepted that the optically-active Car S_2_ state is involved in light harvesting, with excitation energy transfer from Car S_2_ to Chl Q_x_ (or Car S_2_ → Car S_1_ followed by Car S_1_ → Chl Q_y_), while quenching may involve the optically-dark S_1_ or one of the theorised, yet poorly characterised, intermediate excited states of Cars (Liguori et al., 2017). Likewise, there have been suggestions that an intermediate Lut excited state is also involved in light harvesting (J. Li et al., 2023; Ma et al., 2025; Macernis et al., 2012; Son et al., 2019). Whilst the excitonic character of these Lut states is debated, it is generally agreed that pigment conformational change can impact the excitation levels of Lut, including these possible intermediate states. The nature and degree of conformational change of the Lut have been considered in investigations that employed molecular dynamics simulations (Pedraza-González et al., 2024), and the effect has been modelled by quantum chemical calculations, starting with the previously solved LHCII structural models (Macernis et al., 2012; T. Wei et al., 2019). Specifically, twists within the backbone and end rings of Lut have been found to impact its energy levels and, whilst some studies have suggested this is not enough alone to induce a switch from light harvesting to quenching (Macernis et al., 2012; T. Wei et al., 2019), other studies have proposed that it could be (Ilioaia et al., 2013; Liguori et al., 2017; Ma et al., 2025; Son et al., 2019). We may have pinpointed some rotational changes of importance for Lut1/2 in our new LHCII model (Fig. 5, Table 2) that we highlight below.

To try to understand the difference between a light harvesting and quenching state of Lut, again, comparisons of our new LHCII models to the crystal structure model were made. The crystal structure has a slightly lower resolution of the Lut than our models, visible as a slightly more diffuse electron density in the X-ray map (Fig. 6C, green map) and therefore, there is some doubt in the exact conformations of these pigments. When measuring the rotation of the Lut backbones in our LHCII model (at pH 7.5) we observe a 17-18° rotation in Lut1 on both stromal and lumenal sides (Fig. 5 and Table 2, bold). Meanwhile, Lut1 within the crystal structure is straighter, with 9° rotation on the lumenal side and 4° on the stromal side (Fig. 6 and Table 2). However, when assessing the rotation of the end rings, the Lut1 in our LHCII model has a lesser twist than the crystal structure whether considering the lumenal end (64° vs. 77°) or the stromal end (117° vs. 121°), as seen in Table 2 and Fig. 6C. Whilst Lut1 is thought to be the main quencher, Lut2 should also be considered. Again, comparing our LHCII model (pH 7.5) to the one from crystal structures, Lut2 has significant differences in the backbone and end-group rotation (Table 2).

The nature of the distortions that we observe (Car backbone and end-ring rotations) and the approximate amplitude of our distortions, do correlate with some previously completed excitonic calculations (Ma et al., 2025; T. Wei et al., 2019), suggesting that they could have a significant impact on the excitation dynamics within LHCII. It may be interesting to revisit previous studies of excitation energy transfer and dissipation in view of these new pigment conformations, for example, the quantum mechanical modelling and excited state calculations of Fox et al. (2015) predict an even twisting along the Lut2 backbone but Lut1 rotation localised to one end. Given the range of conflicting reports in the literature regarding Car energetics, it is feasible that the Lut within LHCII takes on a range of conformations which allow for a number of different energy levels and intermediates, consistent with the dynamic equilibrium of energetic states found in single-protein fluorescence spectroscopy of isolated LHCII (Schlau-Cohen et al., 2015; Schörner et al., 2015). We hope that our high resolution and clear Lut conformational information can inform future studies to clarify the relationship between the structure and photophysics of LHCII.

## Conclusion

We have presented three new, high resolution, maps of LHCII in detergent representing two conformations, one shared by samples at both pH 7.5 and 4.5 and one unique to the low pH environment. The high quality of the maps reveals the different conformations of Lut1 and Lut2 which may facilitate new insights into LHCII function. Our comparison of the new LHCII models with the previously published LHCII crystal structures highlights the potential importance of subtle changes in Lut1 conformation in determining the quenching state of LHCII and the invariability of the polypeptide. Future analyses could also make measurements of LHCII helix angles, inter-pigment distances and Lut conformations for the trimeric LHCII that is found within Photosystem II supercomplexes, such as the single-particle cryoEM models of PSII dimers with either two, four or six associated LHCII trimers (Arshad et al., 2026; Shen et al., 2019; X. Wei et al., 2016), where it would be interesting to assess how structural changes relate to their optical properties, although this is complicated by the presence of reaction centres where photochemistry occurs. It would be interesting for theoretical modelling of excited states and energy transfers to be performed using our new structural models.

## Acknowledgements

T.S. was supported by a PhD studentship from the Biotechnology and Biological Sciences Research Council (BBSRC, UK), as part of the Yorkshire Bioscience DTP. P.G.A. was supported by a grant from the BBSRC (BB/W004593/1). The Astbury Biostructure facility, containing the electron microscopy instrumentation used in this work, was funded through the University of Leeds and the Wellcome Trust (198466/Z/15/Z). We would like to thank the staff at the Astbury Biostructure Laboratory at University of Leeds for their excellent support. We also thank Ash Hancock for training T.S. in the biochemical purification of LHCII and Jonathan Machin for writing code to allow T.S. to make measurements in ChimeraX.

