## Supplementary Information for "High resolution structure of plant Light-Harvesting Complex II (LHCII) provides insight into lutein conformations and energy quenching"

### for:

#### Supplementary Figures

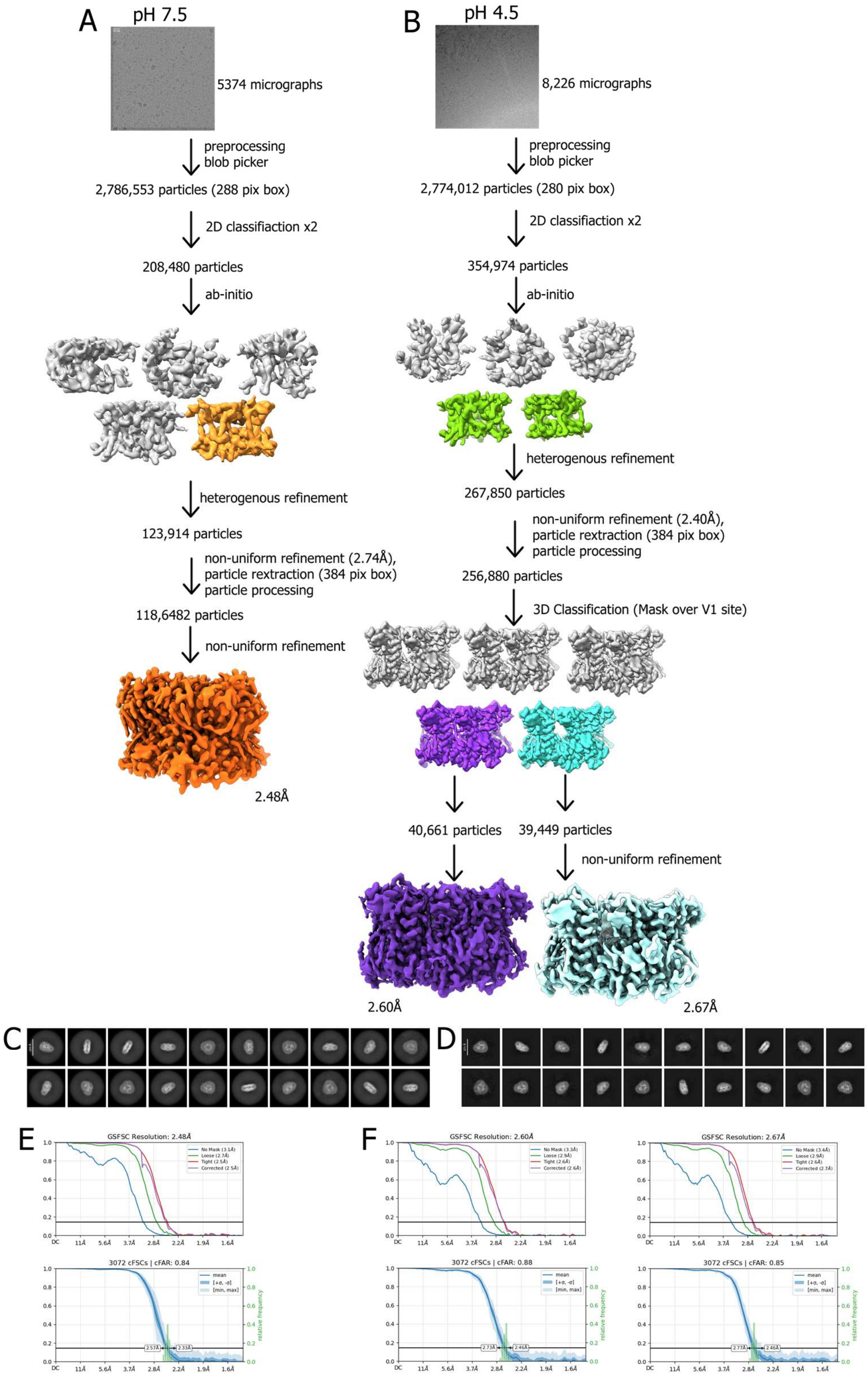


***Fig. S1*** *CryoEM data processing workflow. (**A**) CryoSparc workflow for the pH 7.5 data set to achieve one model at 4.48Å. 3-D classification was attempted with this data set but yielded no alternative structures. (**B**) CryoSparc workflow for the pH 4.5 data set to achieve two final models at 2.60 Å (purple) and 2.67 Å (blue). (**C**) Example 2-D classes for the pH 7.5 data set. (**D**) Example 2-D classes for the pH 4.5 data set. (**E**) FSC curve and cFAR score for pH 7.5 map. (**F**) FSC curve and cFAR score for the two pH 4.5 maps (pH 4.5 model left, destabilised pH 4.5 model right). Coloured models indicate those used in the subsequent steps*

A

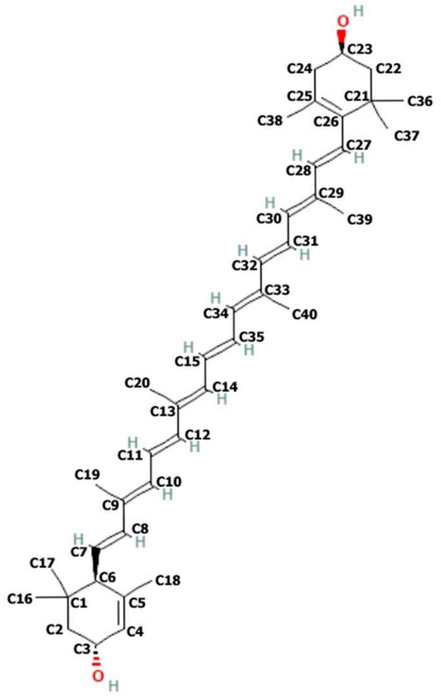


B

| Measurement | Definition |
|---|---|
| RMSD-αC | Root mean square deviation of all polypeptide alpha carbon atoms of two models |
| RMSD-Mg | Root mean square deviation of all chlorophyll Mg atoms between two models for all three trimers |
| Helix E-D separation | Distance between the alpha carbons of V96 and G204 of the same monomer |
| Helix A/B crossing angle | Angle formed by the amino acids F58, C69, R185 and V200 where C69 and R185 forms a pivot point |
| Lutein 1 to Chl α 612 separation | Distance between C15 of Lut1 and the Mg atom of Chl α 612 of the same monomer |
| Lutein 1 to Chl α 610 separation | Distance between C26 of Lut1 and the Mg atom of Chl α 610 of the same monomer |
| Lutein 1 to Chl α 613 separation | Distance between C4 of Lut1 and the Mg atom of Chl α 613 of the same monomer |
| Region II rotation | Torsion measurement C19, C9, C13, C20 of the specified lutein |
| Region III rotation | Torsion measurement C20, C13, C33, C40 of the specified lutein |
| Region IV rotation | Torsion measurement C40, C33, C29, C39 of the specified lutein |
| Region I end ring rotation | Torsion measurement C1, C6, C7, C8 of the specified lutein |
| Region V end ring rotation | Torsion measurement C25, C26, C27, C28 of the specified lutein |

***Fig. S2*** *Definition of measurements. (**A**) Carbon numbering of lutein. (**B**) The measurements made in this paper and how they are defined*

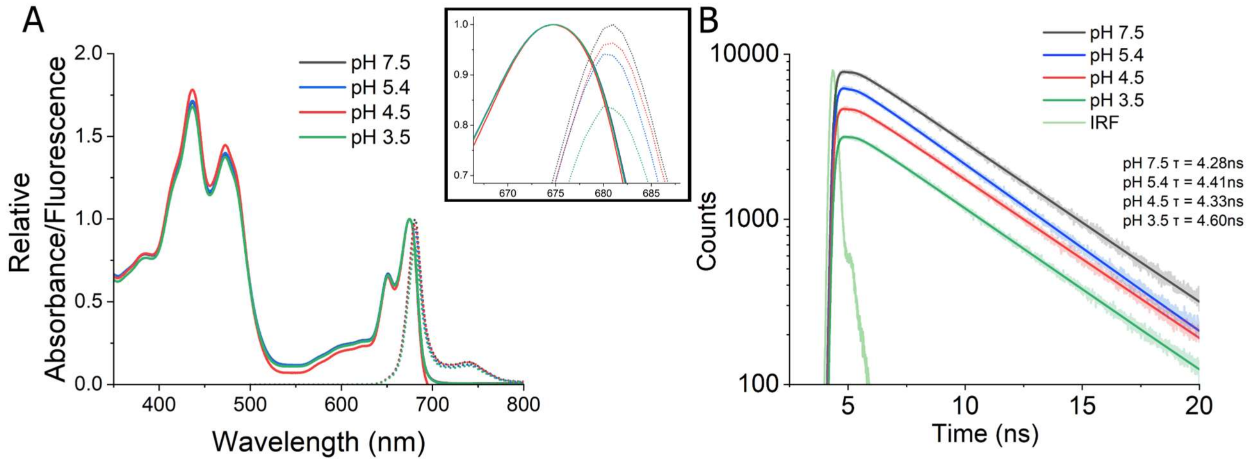


***Fig. S3*** *Absorbance and fluorescence data for LHCII in α-DDM at a range of pH. (**A**) Absorbance (solid line) and emission spectra (dotted line, 436nm excitation) of LHCII at pH 7.5 (black) and pH 4.5 (red). The absorbance is normalised to 1.0 at 675 nm. The fluorescence emission peak was divided by the absorbance at 436 nm for the sample to account for any difference in concentration. Then the peak maximum was corrected to 1.0 for the LHCII pH 7.5 sample by dividing by the counts at peak maximum and the other fluorescence spectra were divided by the same factor (so that all peak heights are relative to the LHCII pH 7.5 sample). (**B**) Fluorescence decay curves and fits for LHCII samples at pH 3.5, 4.5, 5.4 and 7.5. The mean fluorescence lifetimes calculated from an exponential fit are 4.60 ns 4.33 ns, 4.41 ns and 4.28 ns respectively. Excitation was provided by a 475 nm laser and photons were collected at 680 nm. Fitted curves are displayed as solid lines, raw data as transparent lines. The instrument response function (IRF) is green. The curves are offset from each other, by translating vertically by an arbitrary amount, for visibility. Graphs created with Origin 2025*

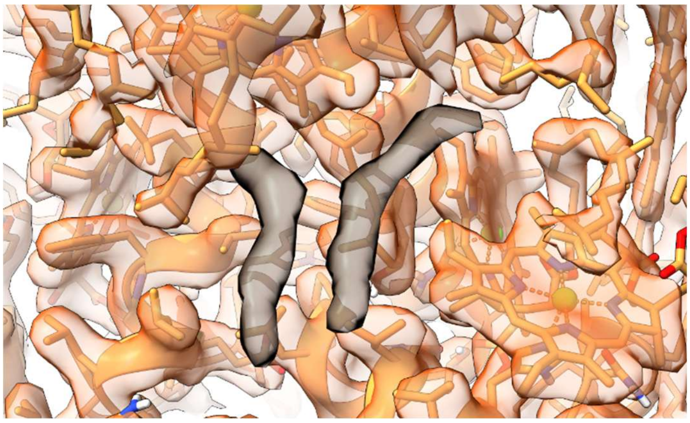

***Fig. S4*** *Unidentified electron density. CryoEM map of LHCII at pH 7.5 with atomic model. The unmodelled density is shown in black. It is likely to be detergent or lipid, but the identity is unclear. The same density is present at pH 4.5*

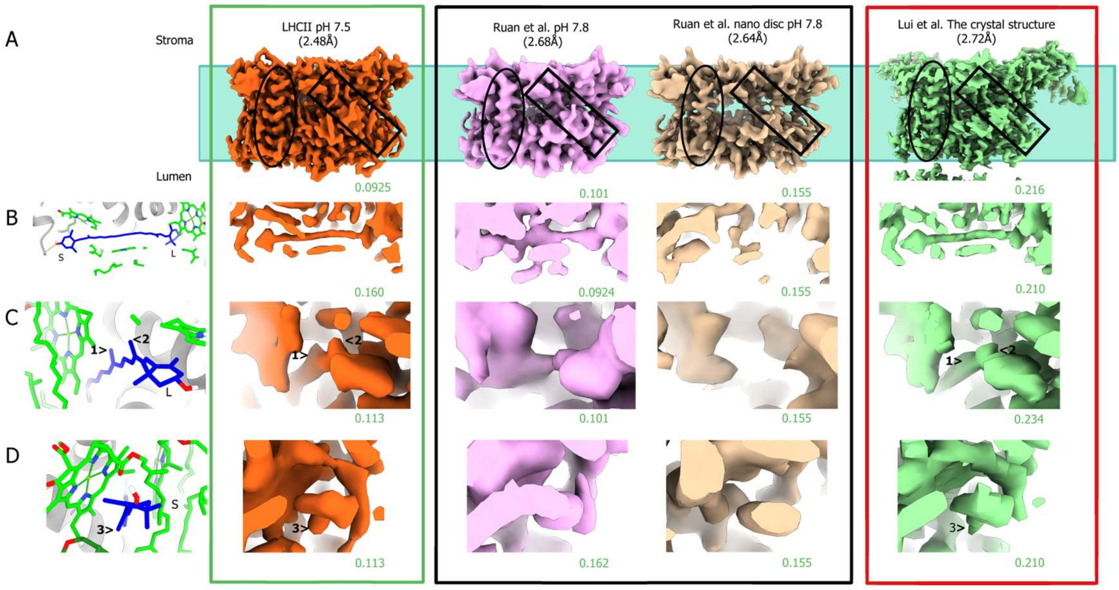


***Fig. S5*** *Comparison of map density for LHCII at neutral pH. (**A**) Side view of each map, aligned as if looking through the membrane, with the approximate lipid membrane location shown by the blue box. The origin (publication) of the map is labelled (above) with the reported resolution shown in brackets. Helix E is circled as a useful point of comparison for map quality. There is more definition in the LHCII pH 7.5 and crystal structure maps than either of the LHCII maps from Ruan et al. The location of lutein 1 is shown by a box, though it sits behind a chlorophyll when viewed from this angle. (**B**) View of lutein 1 for each map. The model of this region from our pH 7.5 model is shown on the far left, lumenal side labelled L, Stromal side labelled S. The shape of the lutein is well defined for our map of LHCII at pH 7.5 and the crystal structure. The Ruan et al. pH 7.8 map is ill-defined with the electron density merging into surrounding regions so there is significant uncertainty to any molecules fitted. The Ruan et al. nanodisc pH 7.8 map has little visible density for lutein at this contour, however, reducing the contour further introduces significant noise from lipids. (**C**) View of lutein 1 from the lumenal side. The head shape is clear in our LHCII pH 7.5 map with side carbons (1,2) clearly visible, so the molecule's orientation and conformation are nicely restrained by the map. The side carbons are also somewhat visible in the crystal structure map but missing from the Ruan et al. maps, so lutein could have various conformations or orientations allowed by the map. (**D**) Lutein 1 viewed from the stromal side. The shape of the lutein is well defined in our LHCII pH 7.5 map with the side carbon (3) clearly visible. The shape is less clear in the crystal structure but still recognisable, whereas, the Ruan et al. maps have only a round shape present so any molecules fitted would have few restraints. For each image the contour levels are shown in green below and the overall reported map resolution is shown above in brackets. Contour levels were chosen as they demonstrate the region best without the introduction of lipid noise to the map. Our cryoEM map (green box): LHCII at pH 7.5 (orange). Ruan et al. 2023 cryoEM maps (black box): LHCII at pH 7.8 in detergent, not protonated (pink) and LHCII at pH 7.8 in nanodisc, not protonated (beige). Lui et al 2004 (red box): the crystal structure X-ray diffraction map (green)*

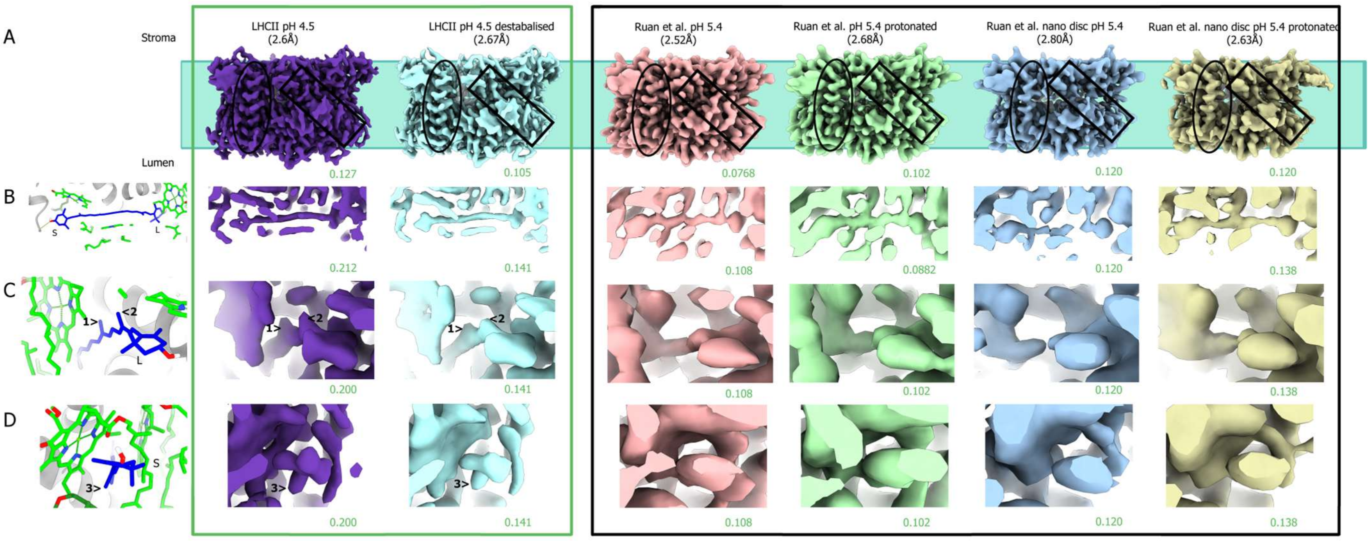


***Fig. S6*** *Comparison of map density for LHCII at low pH. The labelling is the same as in Fig. S5. In both our work (green box) and Ruan et al. (black box), two conformations are found at lowered pH for each detergent or nanodisc dataset.*

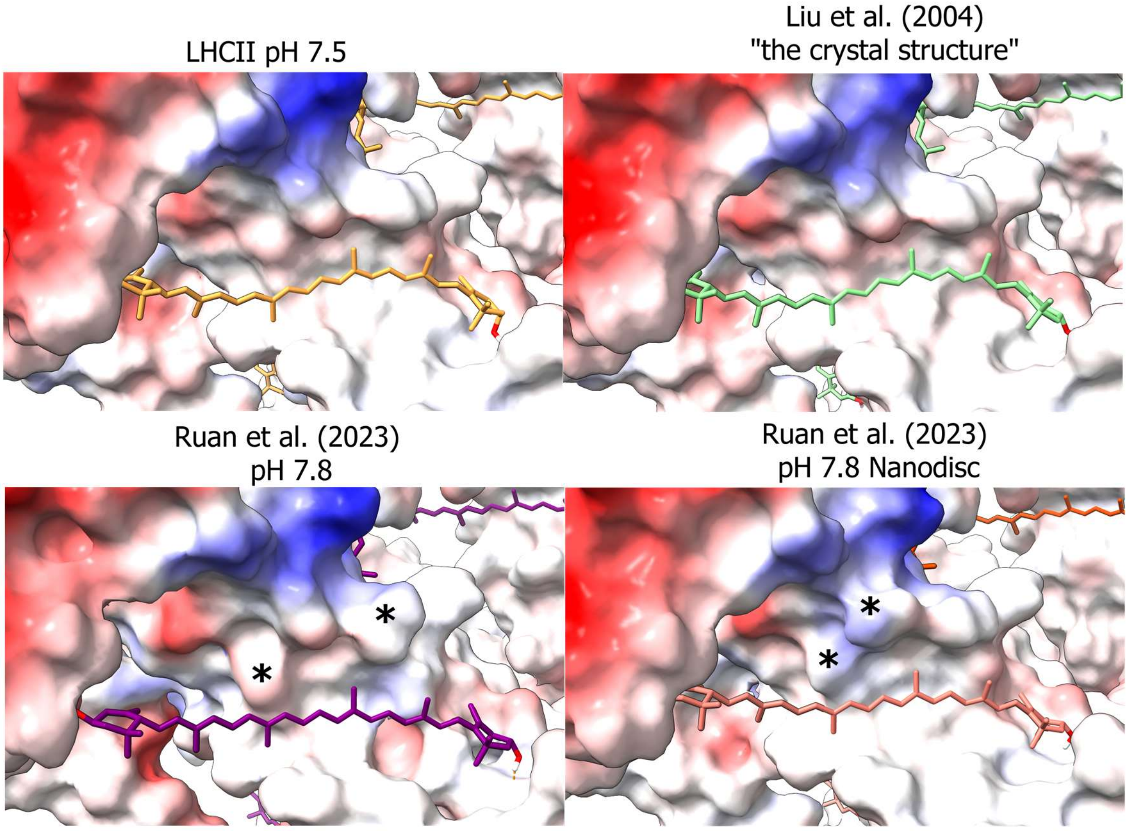


***Fig. S7*** *Comparison of Lut1 binding pockets. Coulombic electrostatic potential of the polypeptide of each model was calculated with ChimeraX to visualise the binding pocket of Lut1: red is more negative, blue is more positive. A Lut1 of each model is shown. Our LHCII pH 7.5 model is almost identical to the crystal structure. The models by Ruan et al. have small differences in the surface of the protein (*) and the lutein pigment clearly has a more planar configuration (straighter)*

#### Supplementary Tables

***Table S1*** *CryoEM data collection and processing information*

| | pH 7.5 | pH 4.5 |
|---|---|---|
| **Data Collection** | | |
| Magnification | 165,000 | 165,000 |
| Voltage (kV) | 300 | 300 |
| Electron exposure ($e^-/Å^2$) | 50 | 50 |
| Defocus range (μm) | 0.7-2.7 | 0.7-2.7 |
| Pixel size (Å) | 0.74 | 0.74 |
| **Processing** | | |
| Micrographs | 5374 | 8226 |
| Final extraction box size | 384 | 384 |
| Number of models | 1 | 2 |
| Symmetry imposed | C1 | C1 |

***Table S2*** *Table of Measurements of all models*

| Model | RMSD-αC to our pH 7.5 model | RMSD-Mg to our pH 7.5 model | Separation Helix D-E (Å) | | | | | Helix crossing angle (°) | | | | | Lut1 to Chl *a* 612 (Å) | | | | |
|---|---|---|---|---|---|---|---|---|---|---|---|---|---|---|---|---|---|
| | | | A | B | C | **Mean** | Std. Dev | A | B | C | **Mean** | Std. Dev | A | B | C | **Mean** | Std. Dev |
| pH 7.5 | 0.00 | 0.00 | 5.34 | 5.35 | 5.30 | **5.33** | 0.02 | 117.65 | 117.54 | 118.18 | **117.79** | 0.28 | 5.63 | 5.53 | 5.41 | **5.53** | 0.09 |
| pH 4.5 | 0.24 | 0.49 | 5.29 | 5.43 | 5.39 | **5.37** | 0.06 | 118.04 | 117.60 | 118.28 | **117.98** | 0.28 | 5.50 | 5.50 | 5.41 | **5.47** | 0.04 |
| Destabalised pH 4.5 | 0.29 | 0.54 | 5.25 | 5.34 | 5.26 | **5.28** | 0.04 | 118.57 | 117.22 | 119.42 | **118.40** | 0.91 | 5.31 | 5.30 | 5.26 | **5.29** | 0.02 |
| Liu et al. 07 | 0.31 | 0.44 | 5.45 | 5.42 | 5.53 | **5.46** | 0.05 | 118.52 | 118.28 | 118.46 | **118.42** | 0.10 | 5.51 | 5.60 | 5.59 | **5.57** | 0.04 |
| Standfuss et al. 05 | 0.38 | 0.41 | 5.45 | 5.36 | 5.46 | **5.42** | 0.04 | 117.14 | 117.26 | 117.28 | **117.23** | 0.06 | 5.39 | 5.43 | 5.46 | **5.43** | 0.03 |
| Wan et al. 14 | 0.33 | * | 5.25 | 5.40 | 5.25 | **5.30** | 0.07 | 118.97 | 119.21 | 119.03 | **119.07** | 0.10 | 5.50 | 5.55 | 5.53 | **5.53** | 0.02 |
| Seki et al. 24 (C3) | 0.28 | 0.58 | 5.72 | 5.73 | 5.76 | **5.74** | 0.02 | 119.25 | 119.25 | 119.23 | **119.24** | 0.01 | 5.52 | 5.57 | 5.55 | **5.55** | 0.02 |
| Ruan et al. 23: Detergent pH 7.8 | 0.49 | 0.67 | 5.92 | 5.95 | 5.98 | **5.95** | 0.02 | 120.61 | 120.49 | 121.27 | **120.79** | 0.34 | 6.00 | 6.02 | 5.96 | **5.99** | 0.02 |
| stan | 0.38 | 0.41 | 5.45 | 5.36 | 5.46 | **5.42** | 0.04 | 117.14 | 117.26 | 117.28 | **117.23** | 0.05 | 5.39 | 5.43 | 5.46 | **5.43** | 0.03 |
| wan | 0.33 | | 5.25 | 5.40 | 5.25 | **5.30** | 0.06 | 118.97 | 119.21 | 119.03 | **119.07** | 0.09 | 5.50 | 5.55 | 5.53 | **5.53** | 0.02 |
| Detergent pH 5.4 | 0.54 | 0.74 | 6.01 | 5.97 | 5.99 | **5.99** | 0.01 | 121.72 | 120.78 | 121.10 | **121.20** | 0.39 | 6.10 | 6.05 | 6.15 | **6.10** | 0.04 |
| Detergent pH 5.4 protonated | 1.37 | 1.29 | 6.01 | 6.38 | 6.03 | **6.14** | 0.17 | 118.75 | 116.68 | 118.44 | **117.96** | 0.91 | 5.23 | 6.61 | 6.17 | **6.00** | 0.58 |
| Nanodisc pH 7.8 | 1.46 | * | 7.03 | 7.08 | 6.92 | **7.01** | 0.07 | 121.07 | 119.85 | 121.69 | **120.87** | 0.76 | 5.76 | 5.57 | 5.57 | **5.63** | 0.09 |
| Nanodisc pH 5.4 | 1.72 | 2.09 | 7.15 | 7.16 | 6.94 | **7.09** | 0.10 | 121.29 | 119.65 | 121.73 | **120.89** | 0.90 | 6.28 | 6.08 | 6.27 | **6.21** | 0.09 |
| Nanodisc pH 5.4 protonated | 0.86 | * | 6.46 | 6.50 | 6.19 | **6.38** | 0.14 | 120.97 | 118.29 | 119.12 | **119.46** | 1.12 | 5.91 | 5.39 | 5.47 | **5.59** | 0.23 |

| Model | Lut1 to Chl *a* 610 (Å) | | | | | Lut1 to Chl *a* 613 (Å) | | | | | Lut1 twist region II (°) | | | | |
|---|---|---|---|---|---|---|---|---|---|---|---|---|---|---|---|
| | A | B | C | **Mean** | Std. Dev | A | B | C | **Mean** | Std. Dev | A | B | C | **Mean** | Std. Dev |
| pH 7.5 | 5.54 | 5.67 | 5.57 | **5.60** | 0.05 | 4.92 | 4.81 | 4.77 | **4.83** | 0.06 | 17.4 | 19.8 | 16.4 | **17.9** | 1.4 |
| pH 4.5 | 5.64 | 5.64 | 5.66 | **5.65** | 0.01 | 4.53 | 4.59 | 4.65 | **4.59** | 0.05 | 11.9 | 17.6 | 19.0 | **16.2** | 3.1 |
| Destabalised pH 4.5 | 5.39 | 5.59 | 5.53 | **5.50** | 0.09 | 5.40 | 4.72 | 4.77 | **4.96** | 0.31 | 13.3 | 6.9 | 13.3 | **11.2** | 3.0 |
| Liu et al. 07 | 5.81 | 5.72 | 5.66 | **5.73** | 0.06 | 4.98 | 4.95 | 4.86 | **4.93** | 0.05 | 8.7 | 9.0 | 9.0 | **8.9** | 0.2 |
| Standfuss et al. 05 | 5.71 | 5.70 | 5.68 | **5.70** | 0.01 | 5.10 | 5.15 | 5.09 | **5.11** | 0.03 | 1.1 | 1.9 | 1.2 | **1.4** | 0.4 |
| Wan et al. 14 | 5.75 | 5.73 | 5.54 | **5.67** | 0.09 | 5.06 | 5.18 | 4.92 | **5.05** | 0.10 | 13.1 | 12.5 | 18.2 | **14.6** | 2.6 |
| Seki et al. 24 (C3) | 5.626 | 5.599 | 5.635 | **5.62** | 0.02 | 4.559 | 4.611 | 4.627 | **4.60** | 0.03 | 13.7 | 12.5 | 13.2 | **13.1** | 0.5 |
| Ruan et al. 23: Detergent pH 7.8 | 5.32 | 5.30 | 5.30 | **5.31** | 0.01 | 4.93 | 4.63 | 4.65 | **4.73** | 0.14 | 22.4 | 23.6 | 27.2 | **24.4** | 2.1 |
| stan | 5.71 | 5.70 | 5.68 | **5.70** | 0.01 | 5.10 | 5.15 | 5.09 | **5.11** | 0.02 | 1.1 | 1.9 | 1.2 | **1.4** | 0.3 |
| wan | 5.75 | 5.73 | 5.54 | **5.67** | 0.08 | 5.06 | 5.18 | 4.92 | **5.05** | 0.09 | 13.1 | 12.5 | 18.2 | **14.6** | 2.2 |
| Detergent pH 5.4 | 5.20 | 5.02 | 5.37 | **5.19** | 0.14 | 5.15 | 4.89 | 4.88 | **4.97** | 0.12 | 22.7 | 22.7 | 30.7 | **25.4** | 3.8 |
| Detergent pH 5.4 protonated | 5.79 | 5.68 | 5.54 | **5.67** | 0.10 | 5.05 | 4.97 | 5.15 | **5.05** | 0.07 | 14.0 | 22.8 | 44.5 | **27.1** | 12.8 |
| Nanodisc pH 7.8 | 5.93 | 5.87 | 6.11 | **5.97** | 0.10 | 6.89 | 6.66 | 6.76 | **6.77** | 0.09 | 27.2 | 29.3 | 26.8 | **27.8** | 1.1 |
| Nanodisc pH 5.4 | 5.95 | 5.62 | 5.91 | **5.83** | 0.15 | 6.69 | 6.29 | 6.13 | **6.37** | 0.23 | 23.2 | 25.5 | 19.9 | **22.8** | 2.3 |
| Nanodisc pH 5.4 protonated | 5.78 | 6.01 | 6.58 | **6.12** | 0.34 | 5.50 | 6.37 | 5.07 | **5.65** | 0.54 | 24.6 | 24.6 | 24.6 | **24.6** | 0.0 |

| Model | Lut1 twist region III (°) | | | | | Lut1 twist region IV (°) | | | | | Lut2 twist region II (°) | | | | |
|---|---|---|---|---|---|---|---|---|---|---|---|---|---|---|---|
| | A | B | C | **Mean** | Std. Dev | A | B | C | **Mean** | Std. Dev | A | B | C | **Mean** | Std. Dev |
| pH 7.5 | 174.7 | 170.1 | 170.1 | **171.6** | 2.2 | 15.4 | 15.8 | 18.4 | **16.5** | 1.4 | 2.9 | 10.3 | 3.3 | **5.5** | 3.4 |
| pH 4.5 | 174.9 | 170.1 | 168.0 | **171.0** | 2.9 | 16.5 | 16.2 | 20.1 | **17.6** | 1.8 | 5.0 | 2.0 | 2.2 | **3.0** | 1.4 |
| Destabalised pH 4.5 | 163.9 | 174.7 | 171.7 | **170.1** | 4.6 | 9.0 | 6.3 | 20.1 | **11.8** | 5.9 | 8.4 | 5.4 | 9.0 | **7.6** | 1.6 |
| Liu et al. 07 | 176.9 | 176.2 | 178.2 | **177.1** | 0.8 | 4.4 | 4.5 | 2.9 | **3.9** | 0.7 | 3.6 | 0.8 | 0.4 | **1.6** | 1.4 |
| Standfuss et al. 05 | 176.5 | 175.9 | 177.9 | **176.8** | 0.8 | 2.5 | 2.5 | 3.5 | **2.8** | 0.5 | 33.0 | 31.7 | 34.8 | **33.2** | 1.3 |
| Wan et al. 14 | 175.8 | 177.8 | 165.1 | **172.9** | 5.6 | 10.6 | 0.7 | 12.8 | **8.0** | 5.3 | 5.2 | 7.0 | 5.5 | **5.9** | 0.8 |
| Seki et al. 24 (C3) | 175.8 | 176.3 | 175.3 | **175.8** | 0.4 | 7.7 | 7.5 | 6.3 | **7.2** | 0.6 | 1.1 | 1.8 | 1.2 | **1.4** | 0.3 |
| Ruan et al. 23: Detergent pH 7.8 | 174.9 | 170.7 | 164.3 | **170.0** | 4.4 | 25.1 | 23.7 | 19.7 | **22.8** | 2.3 | 8.8 | 6.5 | 6.1 | **7.1** | 1.2 |
| stan | 176.5 | 175.9 | 177.9 | **176.8** | 0.7 | 2.5 | 2.5 | 3.5 | **2.8** | 0.4 | 33.0 | 31.7 | 34.8 | **33.2** | 1.1 |
| wan | 175.8 | 177.8 | 165.1 | **172.9** | 4.8 | 10.6 | 0.7 | 12.8 | **8.0** | 4.5 | 5.2 | 7.0 | 5.5 | **5.9** | 0.7 |
| Detergent pH 5.4 | 168.1 | 168.7 | 161.7 | **166.2** | 3.2 | 23.9 | 26.4 | 23.1 | **24.4** | 1.4 | 8.5 | 7.4 | 5.3 | **7.1** | 1.3 |
| Detergent pH 5.4 protonated | 163.3 | 155.6 | 150.9 | **156.6** | 5.1 | 5.1 | 5.5 | 2.9 | **4.5** | 1.2 | 15.8 | 12.3 | 0.6 | **9.6** | 6.5 |
| Nanodisc pH 7.8 | 155.9 | 169.9 | 166.4 | **164.1** | 5.9 | 7.1 | 10.8 | 4.6 | **7.5** | 2.6 | 16.0 | 20.4 | 21.9 | **19.4** | 2.5 |
| Nanodisc pH 5.4 | 149.8 | 169.6 | 176.8 | **165.4** | 11.5 | 5.3 | 10.2 | 3.1 | **6.2** | 3.0 | 9.8 | 15.6 | 22.9 | **16.1** | 5.3 |
| Nanodisc pH 5.4 protonated | 154.6 | 154.6 | 154.6 | **154.6** | 0.0 | 1.9 | 1.9 | 1.8 | **1.9** | 0.0 | 15.0 | 10.9 | 13.2 | **13.0** | 1.7 |

| Model | Lut2 twist region III (°) | | | | | Lut2 twist region IV (°) | | | | | Lut1 ring rotation region I (°) | | | | |
|---|---|---|---|---|---|---|---|---|---|---|---|---|---|---|---|
| | A | B | C | **Mean** | Std. Dev | A | B | C | **Mean** | Std. Dev | A | B | C | **Mean** | Std. Dev |
| pH 7.5 | 170.5 | 179.5 | 173.3 | **174.4** | 3.74 | 21.7 | 24.0 | 28.2 | **24.6** | 2.7 | 63.77 | 67.28 | 61.37 | **64.14** | 2.43 |
| pH 4.5 | 172.3 | 164.7 | 178.9 | **172.0** | 5.83 | 32.2 | 21.4 | 32.1 | **28.6** | 5.1 | 66.97 | 71.42 | 63.57 | **67.32** | 3.22 |
| Destabalised pH 4.5 | 174.7 | 175.0 | 175.0 | **174.9** | 0.12 | 33.7 | 20.9 | 27.4 | **27.4** | 5.2 | 99.32 | 97.82 | 98.97 | **98.70** | 0.64 |
| Liu et al. 07 | 171.8 | 170.3 | 174.3 | **172.1** | 1.65 | 10.9 | 11.2 | 10.3 | **10.8** | 0.4 | 77.00 | 77.41 | 76.70 | **77.04** | 0.29 |
| Standfuss et al. 05 | 172.6 | 173.0 | 172.1 | **172.5** | 0.38 | 8.3 | 6.5 | 9.0 | **8.0** | 1.1 | 94.81 | 98.73 | 96.55 | **96.70** | 1.60 |
| Wan et al. 14 | 160.5 | 155.1 | 152.7 | **156.1** | 3.29 | 16.2 | 22.7 | 25.7 | **21.5** | 4.0 | 85.02 | 78.12 | 85.46 | **82.87** | 3.36 |
| Seki et al. 24 (C3) | 170.0 | 171.3 | 171.9 | **171.1** | 0.79 | 20.7 | 21.0 | 17.7 | **19.8** | 1.5 | 85.431 | 84.7 | 85.08 | **85.06** | 0.31 |
| Ruan et al. 23: Detergent pH 7.8 | 178.0 | 178.7 | 176.9 | **177.9** | 0.77 | 28.1 | 34.3 | 30.0 | **30.8** | 2.6 | 94.24 | 82.77 | 91.22 | **89.41** | 4.85 |
| stan | 172.6 | 173.0 | 172.1 | **172.5** | 0.33 | 8.3 | 6.5 | 9.0 | **8.0** | 0.9 | 94.81 | 98.73 | 96.55 | **96.70** | 1.39 |
| wan | 160.5 | 155.1 | 152.7 | **156.1** | 2.85 | 16.2 | 22.7 | 25.7 | **21.5** | 3.4 | 85.02 | 78.12 | 85.46 | **82.87** | 2.91 |
| Detergent pH 5.4 | 178.8 | 180.0 | 174.3 | **177.7** | 2.48 | 29.7 | 33.6 | 30.1 | **31.1** | 1.7 | 93.86 | 83.08 | 91.82 | **89.59** | 4.68 |
| Detergent pH 5.4 protonated | 174.3 | 174.2 | 163.6 | **170.7** | 5.00 | 19.1 | 42.6 | 27.6 | **29.7** | 9.7 | 70.34 | 78.14 | 85.34 | **77.94** | 6.12 |
| Nanodisc pH 7.8 | 163.0 | 172.0 | 166.4 | **167.1** | 3.74 | 34.6 | 9.2 | 32.5 | **25.4** | 11.5 | 86.42 | 80.95 | 88.12 | **85.16** | 3.06 |
| Nanodisc pH 5.4 | 167.8 | 177.3 | 166.3 | **170.5** | 4.85 | 29.2 | 15.6 | 32.1 | **25.6** | 7.2 | 90.64 | 86.88 | 84.60 | **87.37** | 2.49 |
| Nanodisc pH 5.4 protonated | 174.8 | 174.8 | 171.5 | **173.7** | 1.54 | 31.6 | 33.9 | 27.3 | **30.9** | 2.7 | 44.03 | 44.15 | 44.07 | **44.08** | 0.05 |

| Model | Lut1 ring rotation region V (°) | | | | | Lut2 ring rotation region I (°) | | | | | Lut2 ring rotation region V (°) | | | | |
|---|---|---|---|---|---|---|---|---|---|---|---|---|---|---|---|
| | A | B | C | **Mean** | Std. Dev | A | B | C | **Mean** | Std. Dev | A | B | C | **Mean** | Std. Dev |
| pH 7.5 | 112.1 | 115.8 | 123.7 | **117.2** | 4.8 | 74.5 | 74.5 | 77.0 | **75.3** | 1.2 | 139.5 | 140.4 | 137.7 | **139.2** | 1.1 |
| pH 4.5 | 125.3 | 123.2 | 126.3 | **124.9** | 1.3 | 71.7 | 69.3 | 70.5 | **70.5** | 1.0 | 136.3 | 136.0 | 137.0 | **136.4** | 0.4 |
| Destabalised pH 4.5 | 122.9 | 125.3 | 127.5 | **125.3** | 1.9 | 63.2 | 63.9 | 66.4 | **64.5** | 1.4 | 134.8 | 140.9 | 139.1 | **138.3** | 2.6 |
| Liu et al. 07 | 121.0 | 122.7 | 119.7 | **121.2** | 1.2 | 81.7 | 79.6 | 79.1 | **80.2** | 1.1 | 129.1 | 126.4 | 127.1 | **127.5** | 1.2 |
| Standfuss et al. 05 | 6.5 | 5.3 | 7.9 | **6.6** | 1.1 | 105.3 | 109.1 | 112.2 | **108.9** | 2.8 | 3.8 | -4.0 | 3.4 | **1.1** | 3.6 |
| Wan et al. 14 | 128.2 | 126.1 | 132.7 | **129.0** | 2.8 | 80.1 | 78.4 | 80.7 | **79.7** | 1.0 | 136.0 | 123.9 | 130.8 | **130.3** | 5.0 |
| Seki et al. 24 (C3) | 124.6 | 126.3 | 124.2 | **125.0** | 0.9 | 84.5 | 82.8 | 82.5 | **83.2** | 0.9 | 131.7 | 131.2 | 131.8 | **131.6** | 0.2 |
| Ruan et al. 23: | | | | | | | | | | | | | | | |
| Detergent pH 7.8 | 111.5 | 121.7 | 121.9 | **118.4** | 4.8 | 84.0 | 79.5 | 84.6 | **82.7** | 2.3 | 124.0 | 119.5 | 127.1 | **123.5** | 3.1 |
| stan | 6.5 | 5.3 | 7.9 | **6.6** | 0.9 | 105.3 | 109.1 | 112.2 | **108.9** | 2.4 | 3.8 | -4.0 | 3.4 | **1.1** | 3.1 |
| wan | 128.2 | 126.1 | 132.7 | **129.0** | 2.4 | 80.1 | 78.4 | 80.7 | **79.7** | 0.8 | 136.0 | 123.9 | 130.8 | **130.3** | 4.3 |
| Detergent pH 5.4 | 111.1 | 119.9 | 120.4 | **117.1** | 4.3 | 84.3 | 79.7 | 83.9 | **82.6** | 2.1 | 123.1 | 118.7 | 126.1 | **122.7** | 3.0 |
| Detergent pH 5.4 protonated | 107.6 | 113.0 | 122.7 | **114.4** | 6.2 | 76.2 | 76.8 | 90.3 | **81.1** | 6.5 | 120.2 | 114.6 | 118.6 | **117.8** | 2.4 |
| Nanodisc pH 7.8 | 123.2 | 123.0 | 116.0 | **120.7** | 3.4 | 89.2 | 76.9 | 92.5 | **86.2** | 6.7 | 113.1 | 112.7 | 113.3 | **113.0** | 0.3 |
| Nanodisc pH 5.4 | 123.3 | 118.7 | 119.4 | **120.5** | 2.0 | 95.6 | 79.6 | 70.5 | **81.9** | 10.4 | 111.6 | 111.3 | 113.4 | **112.1** | 0.9 |
| Nanodisc pH 5.4 protonated | 115.5 | 115.5 | 115.5 | **115.5** | 0.0 | 77.3 | 85.5 | 78.4 | **80.4** | 3.6 | 133.4 | 130.0 | 127.3 | **130.2** | 2.5 |

*The individual measurements from each monomer of each model are shown alongside the mean and standard deviation. Our models are highlighted in green, the three crystal structures in orange and the other cryoEM structures in white. The measurements taken from the destabilised pH 4.5 monomer missing the V1 pigment with a disordered C-terminus are underlined (monomer C). Monomer identity (A, B, C) was assigned arbitrarily but is consistent between measurements. Some RMSD-Mg were not taken (*) due to inconsistent naming within the model, making incorrect measurements likely*
